# “Distinct inhibitory neurons differently shape neuronal codes for sound intensity in the auditory cortex”

**DOI:** 10.1101/2023.02.01.526470

**Authors:** Melanie Tobin, Janaki Sheth, Katherine C. Wood, Erin K. Michel, Maria N. Geffen

**Author notes:** Corresponding author: Maria N. Geffen.

## Abstract

Cortical circuits contain multiple types of inhibitory neurons which shape how information is processed within neuronal networks. Here, we asked whether somatostatin-expressing (SST) and vasoactive intestinal peptide-expressing (VIP) inhibitory neurons have distinct effects on population neuronal responses to noise bursts of varying intensities. We optogenetically stimulated SST or VIP neurons while simultaneously measuring the calcium responses of populations of hundreds of neurons in the auditory cortex of male and female awake, head-fixed mice to sounds. Upon SST neuronal activation, noise bursts representations became more discrete for different intensity levels, relying on cell identity rather than strength. By contrast, upon VIP neuronal activation, noise bursts of different intensity level activated overlapping neuronal populations, albeit at different response strengths. At the single-cell level, SST and VIP neuronal activation differentially modulated the response-level curves of monotonic and nonmonotonic neurons. SST neuronal activation effects were consistent with a shift of the neuronal population responses toward a more localist code with different cells responding to sounds of different intensity. By contrast, VIP neuronal activation shifted responses towards a more distributed code, in which sounds of different intensity level are encoded in the relative response of similar populations of cells. These results delineate how distinct inhibitory neurons in the auditory cortex dynamically control cortical population codes. Different inhibitory neuronal populations may be recruited under different behavioral demands, depending on whether categorical or invariant representations are advantageous for the task.

**SIGNIFICANCE:** Information about sounds is represented in the auditory cortex by neuronal population activity that has a characteristic sparse structure. Cortical neuronal populations comprise multiple types of excitatory and inhibitory neurons. Here, we find that activating different types of inhibitory neurons differentially controls population neuronal representations, with one type of inhibitory neurons increasing the differences in the identity of the cells recruited to represent the different sounds, and another inhibitory neuron type changing the relative activity level of overlapping neuronal populations. Such transformations may be beneficial for different types of auditory behaviors, suggesting that these different types of inhibitory neurons may be recruited under different behavioral constraints in optimizing neuronal representations of sounds.

## INTRODUCTION

Sensory cortical neuronal networks are comprised of multiple subtypes of neurons, including excitatory and inhibitory neurons. Inhibitory neurons can be further divided into multiple sub-classes, including somatostatin-expressing (SST) and vasoactive intestinal peptide-expressing (VIP) neurons, which mutually inhibit each other (Campagnola et al., 2022). The activity of these neurons modulates stimulus representations in auditory cortex (AC). Specifically, activating SST neurons reduces and decorrelates cortical activity (Chen et al., 2015), sharpens frequency tuning of AC neurons (Phillips and Hasenstaub, 2016) and contributes to surround suppression (Lakunina et al., 2020) and adaptation to stimulus context (Natan et al., 2015, 2017). By contrast, VIP neurons disinhibit excitatory neurons (Millman et al., 2020), largely via their projections onto SST neurons (Pfeffer et al., 2013; Pi et al., 2013), without affecting frequency tuning (Bigelow et al., 2019) and can enable high-excitability states in the cortex (Jackson et al., 2016). Both SST and VIP neurons can be modulated: by noradrenergic and cholinergic inputs for SST neurons (Kawaguchi and Shindou, 1998; Fanselow et al., 2008; Chen et al., 2015), and multiple neuromodulators for VIP neurons (Fu et al., 2014; Zhang et al., 2014; Chen et al., 2015) and therefore may serve differential modulatory functions in the context of different behavioral demands.

Sound pressure level representation supports sound detection, hearing in noise, source localization and distance to target calculation (Litovsky and Clifton, 1992). Most neurons in AC respond selectively to sounds at different sound pressure levels, either in a monotonic or a non-monotonic fashion. Monotonic neurons increase their firing rate with sound intensity, differing in their threshold and slope of the response functions. Non-monotonic neurons exhibit preference for specific sound pressure level ranges, differing in their preferred sound pressure level (Zhang et al., 2013). Previous work found that a mix of monotonic and non-monotonic neurons in the auditory cortex is important for sound encoding (Sun et al., 2017). Because the excitatory neurons in the cortex form tightly connected circuits with inhibitory neurons, inhibitory neuronal activity can shift the sound level response functions of excitatory neurons across monotonic and non-monotonic neurons. These changes can in turn affect the representation of sound pressure level by cortical populations. Whereas multiple studies have examined the effects of SST and VIP neuronal modulation on sound responses in individual neurons in AC(Natan et al., 2015, 2017; Seybold et al., 2015; Phillips and Hasenstaub, 2016; Bigelow et al., 2019; Millman et al., 2020; Seay et al., 2020), their effect on population representation of sound pressure level remains to be fully understood.

Here, we studied whether and how cortical inhibitory neurons control the representation of sound pressure levels in populations of neurons, by presenting periodic noise bursts at different sound pressure levels to awake head-fixed mice and imaging Calcium responses in populations of hundreds of neurons while simultaneously activating SST or VIP neurons optogenetically (Figure 1B, Figure 2). First, we tested whether and how activation of SST or VIP neurons differentially modulated sound pressure level responses at the level of individual neuronal response functions in AC. Next, we tested how the representation of sound pressure level changed in the neuronal space with and without SST and VIP neuronal activation. We tested for the effects of SST and VIP neuronal activation on response sparseness and separation angle of population response vectors. Finally, we tested whether and how changes in the response-level curves of monotonic and nonmonotonic individual neurons upon SST or VIP neuronal activation mediated the changes in representation at the population for monotonic and nonmonotonic neurons. Our results suggest that SST and VIP neuronal activation differentially affect both monotonic and non-monotonic neuronal sound pressure level response functions, thereby shifting the neuronal population codes between localist and distributed representations.

**FIGURE 1:**
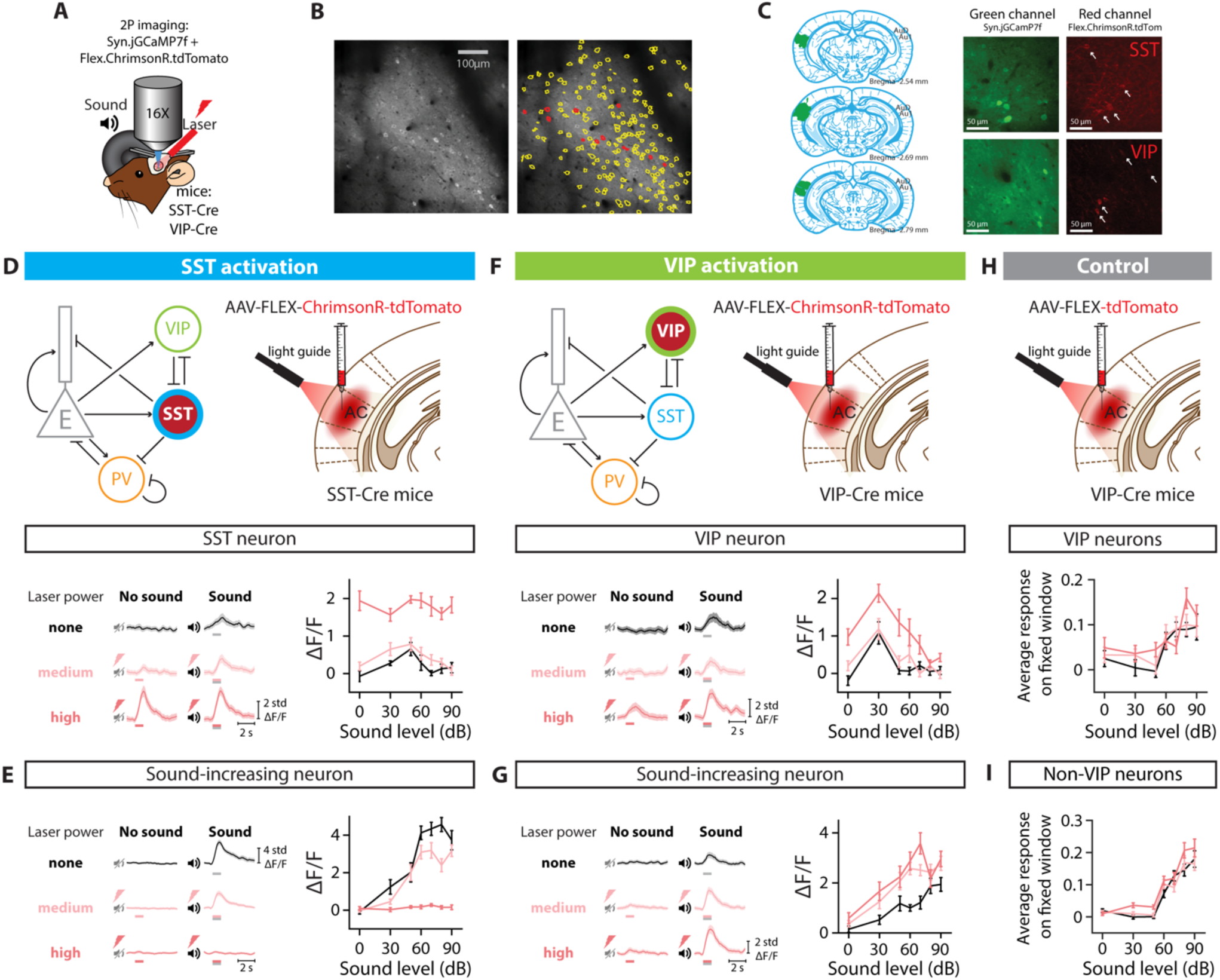
EXPERIMENTAL SET UP AND OPTOGENETIC STIMULATION OF INTERNEURON POPULATIONS. A. Two-photon imaging and laser stimulation through the round window of a mouse injected with viruses encoding Syn.jGCaMP7f and Flex.ChrimsonR.tdTomato in the left Auditory Cortex. Flex.ChrimsonR.tdTomato is injected in AC of SST-Cre and VIP-Cre mice, and is activated by a 635-nm laser. The mouse lines used were SST-Cre x Cdh23+/+ and VIP-Cre x Cdh23+/+. A speaker delivers a broadband noise stimulus at sound pressure levels within 0-90 dB to the right ear. B. Cell tissue with two-photon imaging in the green channel (left) and cell identification (right) using Suite2p software, with yellow lines delineating cell borders, and red lines indicating the neurons expressing ChrimsonR.tdTomato. C. Left: Outline of the spread of the viral injection in a representative brain. Signal in the green channel (center panels) and red channel (right panels) of an SST-Cre mouse (top panels) and an VIP-Cre mouse (bottom panels). Cells identified as VIP or SST interneurons are indicated with an arrow (see Methods). D. (Top) Diagrams for optogenetic manipulation in the circuit and experimental set-up. (Bottom left) Response of a SST neuron to no sound and sound stimulation (50 dB) when activating SST neurons with different laser powers. (Bottom right) Average fluorescence over a 1-s fixed time window (delay from stimulus onset: 90 ms) as a function of sound pressure level for the example cell in the left panel. E. (Left) Response of a sound-increasing neuron to no sound and sound stimulation (70 dB) when activating SST neurons with different laser powers. (Right) Average fluorescence over a 1-s fixed time window (delay from stimulus onset: 150 ms) as a function of sound pressure level for the example cell in the left panel. F. (Top) Diagrams for optogenetic manipulation in the circuit and experimental set-up. (Bottom left) Response of a VIP neuron to no sound and sound stimulation (70 dB) when activating VIP neurons with different laser powers. (Bottom right) Average fluorescence over a 1-s fixed time window (delay from stimulus onset: 300 ms) as a function of sound pressure level for the example cell in the left panel. G. (Left) Response of a sound-increasing neuron to no sound and sound stimulation (70 dB) when activating VIP neurons with different laser powers. (Right) Average fluorescence over a 1-s fixed time window (delay from stimulus onset: 270 ms) as a function of sound pressure level for the example neuron in the left panel. H. (Top) Experimental set-up for the control experiment – laser effect in the absence of an opsin. (Bottom) Average fluorescence over a 1-s fixed window as a function of sound pressure level for the whole population of VIP neurons recorded, tagged with Flex.tdTomato, when the laser illuminates AC. I. Average fluorescence over a 1-s fixed window as a function of sound pressure level for the whole population of neurons recorded (VIP neurons excluded) when the laser illuminates AC. For all panels, black, pink and red colors correspond to no laser power (0 mW/mm2), medium laser power (∼0.3 mW/mm2) and high laser power (∼3.5 mW/mm2), respectively (see Methods for power calibration). The gray and red bars below the example traces in panels D-G indicate the presence of the sound and laser stimulus, respectively.

**FIGURE 2:**
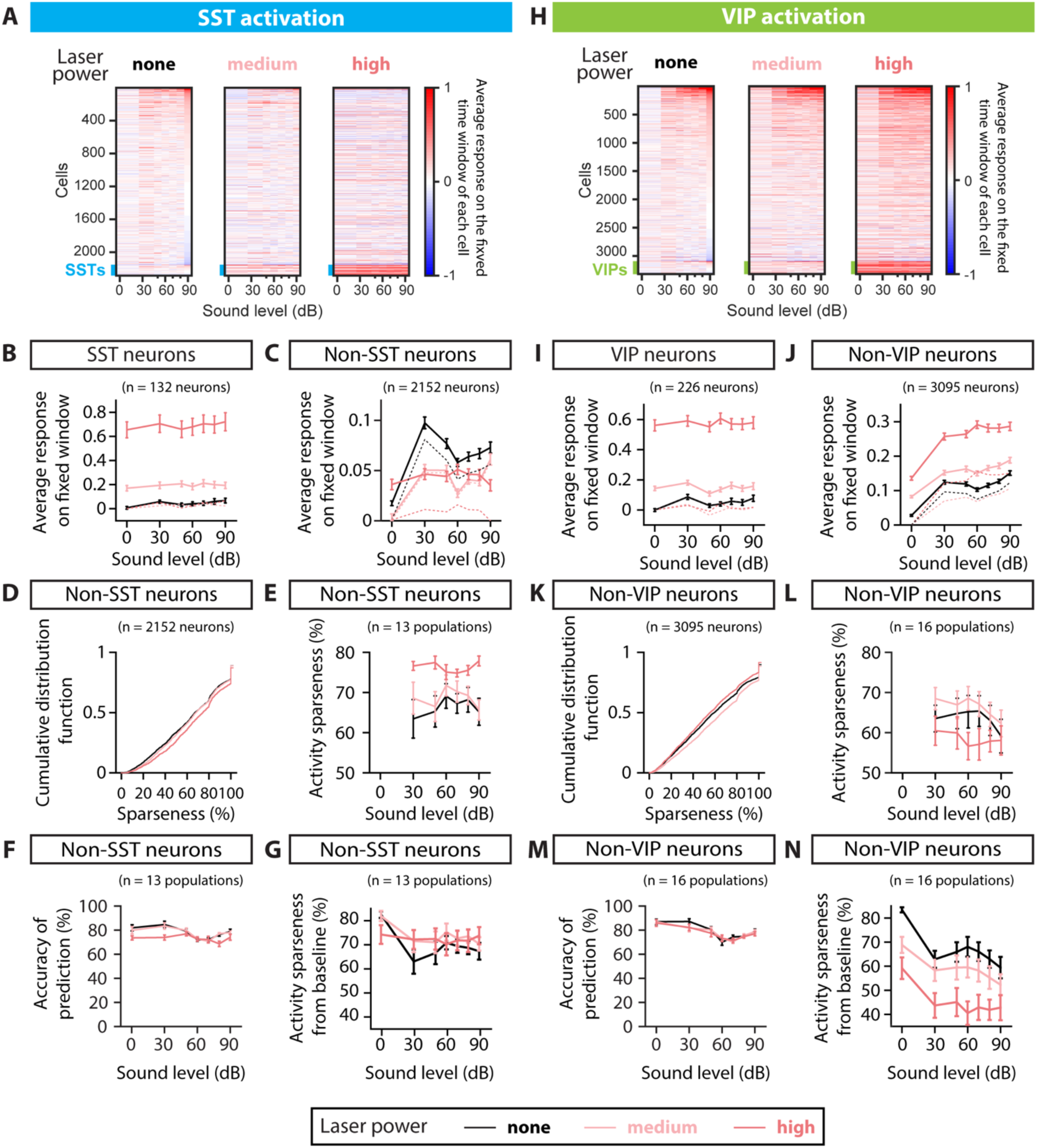
POPULATION RESPONSE TO SST AND VIP NEURONAL ACTIVATION AND SPARSENESS. A. Rasters of the average fluorescence versus sound pressure level for all neurons imaged in the SST-Cre mice, calculated over the 1-s fixed time window (see Methods). Rasters from left to right correspond to SST neuronal activation with no laser power, medium laser power and high laser power. The thick blue line at the bottom of each raster indicates the SST interneurons. Cells are ordered given their response at 90dB and no laser power. B. Absolute average fluorescence (solid lines) and change in average fluorescence relative to the laser and silence condition (dashed lines) over a 1-s fixed window as a function of sound pressure level for the whole population of SST neurons recorded (132 neurons), when the SST neurons are activated. The solid and dashed black line nearly overlap. C. Absolute average fluorescence (solid lines) and change in average fluorescence relative to the laser and silence condition (dashed lines) over a 1-s fixed window as a function of sound pressure level for the whole population of non-SST neurons recorded (2152 neurons), when the SST neurons are activated. D. Cumulative distribution function of sparseness normalized on the population of non-SST neurons, when the SST neurons are activated. Sparseness was defined for non-SST neurons with an increase in response to sound compared to silence at a given laser, corresponding to 2033, 2029 and 1994 neurons for no, mid and high laser, respectively. E. Average activity sparseness measured from silence at a given laser power as a function of sound pressure level for each population of neurons (SST neurons excluded), when the SST neurons are activated. The points at 0dB were by design 100% and thus omitted from the plot and the statistical test. F. Decoding accuracy of SVM decoder at each laser power, decoding individual sound pressure levels using non-SST neuronal responses within 1-s fixed window, when the laser illuminates AC and stimulates SST neurons. G. Average activity sparseness measured baseline as a function of sound pressure level for each population of neurons (SST neurons excluded), when the SST neurons are activated. There was a significant interaction between sound and laser amplitude, but no significant sound-independent laser effect in the activity sparseness measured from baseline upon SST activation (p_laser_=0.37, *p_laser:sound_=4.5e-2, GLME). H. Rasters of the average fluorescence versus the sound pressure level for all neurons imaged in the VIP-Cre mice, calculated over the 1-s fixed time window (see Methods). Rasters from left to right correspond to VIP activation with no laser power, medium laser power and high laser power. The thick green line at the bottom of each raster indicates the VIP interneurons. Cells are ordered given their response at 90dB and no laser power. I. Absolute average fluorescence (solid lines) and change in average fluorescence relative to the laser and silence condition (dashed lines) over a 1-s fixed window as a function of sound pressure level for the whole population of VIP neurons recorded (226 neurons), when the VIP neurons are activated. The solid and dashed black line nearly overlap. J. Absolute average fluorescence (solid lines) and change in average fluorescence relative to the laser and silence condition (dashed lines) over a 1-s fixed window as a function of sound pressure level for the whole population of non-VIP neurons recorded (3095 neurons), when the VIP neurons are activated. K. Cumulative distribution function of sparseness normalized on the population of non-VIP neurons, when the VIP neurons are activated. Sparseness was defined for non-VIP neurons with an increase in response to sound compared to silence at a given laser, corresponding to 2979, 2921 and 3020 neurons for no, mid and high laser, respectively. L. Average activity sparseness measured from silence at a given laser power as a function of sound pressure level for each population of neurons (VIP neurons excluded) when the VIP neurons are activated. The points at 0dB were by design 100% and thus omitted from the plot and the statistical test. M. Decoding accuracy of SVM decoder at each laser power, decoding individual sound pressure levels using non-VIP neuronal responses within 1-s fixed window, when the laser illuminates AC and stimulates VIP neurons. N. Average activity sparseness measured from baseline as a function of sound pressure level for each population of neurons (VIP neurons excluded), when the VIP neurons are activated. There was a significant decrease in activity sparseness measured from baseline upon VIP activation (***p_laser_=1.8e-14, GLME). For all panels, black, pink and red colors correspond to no laser power (0 mW/mm2), medium laser power (∼0.3 mW/mm2) and high laser power (∼3.5 mW/mm2), respectively (see Methods for power calibration).

## METHODS

### Animals

We performed experiments in fourteen adult mice (7 males and 7 females), which were crosses between Cdh23 mice (B6.CAST-Cdh23^Ahl+/^Kjn, JAX: 002756) and Sst-Cre mice (Sst^tm2.1(cre)Zjh^/J, JAX: 013044; n=5 in experimental group) or Vip-IRES-Cre mice (Vip^tm1(cre)Zjh^/J, JAX: 010908; n=4 in experimental group, n=5 in control group) (Table 1). Mice had access to food and water ad libitum and were exposed to light/dark on a reversed 12h cycle at 28°C. Experiments were performed during the animals’ dark cycle. Mice were housed individually after the cranial window implant. All experimental procedures were in accordance with NIH guidelines and approved by the Institutional Animal Care and Use Committee at the University of Pennsylvania.

**Table 1.** Mouse strains and numbers

| Experiment | Figures | Strain | Number of mice | Number of recordings |
| --- | --- | --- | --- | --- |
| GCaMP7f + ChrimsonR | Figs 1-5 | CDH23 x VIP-Cre | 2 | 7 |
|  |  | CDH23 x SST-Cre | 5 | 13 |
| GCaMP6m + ChrimsonR | Figs 1-5 | CDH23 x VIP-Cre | 2 | 9 |
| Control: GCaMP7f + Flex.tdTomato – VIP cells | Fig 1H | CDH23 x VIP-Cre | 4 | 7 |
| Control: GCaMP7f + Flex.tdTomato – Non-VIP cells | Fig 1I | CDH23 x VIP-Cre | 2 | 4 |

### Surgery procedures

Mice were implanted with cranial windows over Auditory Cortex following a published procedure (Wood et al., 2022). Briefly, mice were anesthetized with 1.5-3% isoflurane and the left side of the skull was exposed and perforated by a 3mm biopsy punch over the left Auditory Cortex. We injected in that region 3×750nL of an adeno-associated virus (AAV) mix of AAV1.Syn.GCaMP (6m: Addgene 100841 or 7f: Addgene 104488; dilution 1:10 ∼ 1×10^13^ GC/mL) and AAV1.Syn.Flex.Chrimson.tdTomato (UNC Vector Core; dilution 1:2 ∼ 2×10^12^ GC/mL). In the control mice, we injected a mix of AAV1.Syn.jGCaMP7f (Addgene 104488; dilution 1:10 ∼ 1×10^13^ GC/mL) and AAV1.Syn.Flex.tdTomato (Addgene 28306; dilution 1:100-1:20 ∼ 2×10^11^ – 1×10^12^ GC/mL) in VIP-Cre mice. We then sealed the craniotomy with a glass round window, attached a head plate to the mouse and let it recover for 3-4 weeks. After habituating the mouse to being head fixed for 3 days, we mapped the sound-responsive areas of the brain and located Auditory Cortex using wide field imaging, then performed two-photon imaging in Auditory Cortex (Figure 2C).

### Two-photon imaging

We imaged calcium activity in neurons in layer 2/3 of Auditory Cortex of awake, head-fixed mice (VIP-Cre mice: 3321 neurons over 16 recordings, SST-Cre mice: 2284 neurons over 13 recordings) using the two-photon microscope (Ultima *in vivo* multiphoton microscope, Bruker) with a laser at 940nm (Chameleon Ti-Sapphire). The fluorescence from the tissue went through a Primary Dichroic long pass (620 LP), through an IR Blocker (625 SP), through an Emission Dichroic Long pass (565 LP) which separated the light in two beams. The shorter wavelengths went through an additional bandpass filter (525/70) before being captured by a PMT (“green channel”); the longer wavelengths went through a bandpass filter (595/50) before being captured by a PMT (“red channel”). This set up was used to minimize the contamination of the green channel by the optogenetic stimulus at 635nm. There was nevertheless some bleedthrough during the optogenetic stimulus which was small enough not to saturate the green channel, and thus the activity of neurons could be recorded continuously without interruption during optogenetic stimulation. We checked there was no saturation for the highest laser power used by plotting the average grayscale profile over the dimension of the image perpendicular to the scanning, and verifying it was well below saturation. During a 5-ms laser pulse, while the whole field of view is illuminated by the laser, it appears on the image only for the lines that were being scanned during the laser stimulus. These bands were identified using the average grayscale value of each image line, and the contaminated pixels inside these bands were removed before processing the recordings with Suite2p. We imaged a surface of 512×512 pixels^2^ at 30Hz. If we recorded from a mouse several times, we changed the location or depth within layer 2/3 of Auditory Cortex in order to not image the same neurons twice.

### Optogenetic laser: Power calibration

We first calibrated the laser power by measuring the curve of command voltage versus output power for the laser (Optoengine LLC, MRL-III-635-300mW). The laser’s peak frequency was 635nm. Prior to every recording, we calibrated the laser power at the tissue level as follows: we used an empty cannula to reduce the power of the laser by a factor 10-15 and positioned the optical fiber on the objective so it would shine a spot of 1mm diameter centered on the focal point of the objective. Thus, the calibrated power at the imaging plane was for the medium laser power: 0.3 ± 0.09 mW/mm^2^ (mean ± std, n=29; range: 0.14-0.47 mW/mm^2^) and for the high laser power: 3.4 ± 1.0 mW/mm^2^ (mean ± std, n=29; range: 1.6-5.3 mW/mm^2^).

### Identification of interneurons being stimulated

We started each recording by taking a 2600 frame video both in the green and the red channels (thus imaging GCaMP and tdTomato). As tdTomato is not dependent on the cell activity, any modulation in the signal in the red channel is due to bleedthrough from the GCaMP. We plotted for all cells the raw signal from the red channel versus the signal from the green channel and did a linear fit to extract the bleedthrough coefficient. We then subtracted the bleedthrough in the red signal and calculated the average fluorescence of the processed red signal for every cell. We then z-scored the signal of the red channel to the background fluorescence and selected the cells with a fluorescence higher than 2 σ (standard deviation of the background) as the targeted interneurons. The percentage of cells labeled as VIP or SST interneurons with this criterion was consistent with the percentage of VIP or SST neurons expected within cortex (Rudy et al., 2011).

### Stimulus presentation

We presented combinations of sound and optogenetic stimuli. The auditory stimulus consisted in 1-s long click trains of 25-ms pulses of broadband white noise (range 3−80 kHz) at 10Hz, at 7 sound pressure levels within 0-90 dB SPL (0; 30; 50; 60; 70; 80; 90 dB SPL). The optogenetic stimulus consisted in a 1-s long pulse train of 635nm laser with 5-ms pulses at 20 Hz, at 3 amplitudes with no, medium or high laser power (see power at tissue level in section Optogenetic laser: Power calibration). The two stimuli were presented simultaneously, with the optogenetic stimulus preceding the sound stimulus by 20 ms (Blackwell et al., 2020) for maximal optogenetic effect, the inter-stimulus interval was 5 s. All 21 combinations of sound and optogenetic stimuli were presented randomly and with 10 repeats per combination.

### Analysis of single-cell activity: Fixed time window

For each trial of a stimulus, the response as a function of time was defined as the change in fluorescence ΔF/Fstd compared to the baseline fluorescence Fbaseline over the one-second window preceding the stimulus: ΔF/Fstd = (F – mean(Fbaseline))/std(Fbaseline), with F the fluorescence of the cell as a function of time. *Fixed window:* In order to compare how neuronal responses changed with laser stimulation, keeping all parameters similar besides that one, we defined the fixed window of neuronal response for each recording (one window for all stimulus combinations) as the one-second window with the largest number of responsive neurons. The fixed time window was selected as the 1-s averaging window between [0-1s] and [2-3s] which maximized the number of cells with a significant response to at least one of the stimuli pairs for each recording compared to the pre-stimulus fluorescence (paired t-test, p<0.01 with multiple comparison correction). The delay between the stimulus onset and the beginning of the fixed window was in SST-Cre mice: 385 ± 419 ms (mean ± std, range (min - max): 0 - 1350 ms, n=13) and in VIP-Cre mice: 336 ± 10 ms (mean ± std, range (min - max): 180 - 480 ms, n=16).

Our method for selecting the fixed window gives results similar to a fixed [0 1]s window (Wood et al., 2022), with improvements which we believe have a better chance at capturing the effects of SST or VIP neuronal activation: by allowing for a delay in the beginning of the fixed window, our analysis leads to response-level fits that are closer to the maximum change in fluorescence for most cells.

### Sparseness of a neuron’s response and activity sparseness of a population of neurons

To quantify how many stimuli a neuron responds to, we calculated the sparseness of each neuron adapted from (Vinje and Gallant, 2000):

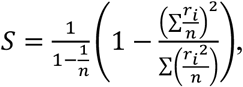

where *r_i_* is the average response of a neuron to the *i*th sound pressure level at a given laser activation calculated over its fixed time window minus the neuron’s response to laser activation and silence, and *n* is the number of sound pressure levels (response in silence excluded). Similar to how this measure is computed with the firing rate of excited neurons (Rolls and Tovee, 1995; Vinje and Gallant, 2000; Olsen and Wilson, 2008; Feigin et al., 2021), we adapted the sparseness measure to fluorescence data by setting any values of *r_i_* < 0 to zero before calculating the sparseness, and by calculating this measure only for neurons with at least one positive *r_i_*, for which the sparseness is well-defined (at least one positive *r_i_*). A sparseness value of 0% indicates that a neuron’s responses to all sound pressure levels are equal, and a sparseness value of 100% indicates that a neuron only responds to one sound pressure level.

To quantify how many neurons in a population are active in response to a given stimulus, we calculated the activity sparseness of each population (Willmore and Tolhurst, 2001) at a given sound level pressure and laser power (Figure 2E and L) by subtracting each neuron’s response over its fixed window to its response at 0dB at the same laser power, and computing the ratio of neurons that had an increase in response above threshold, with the threshold set as described below. The activity sparseness from baseline was computed by taking each neuron’s response over its fixed window, and computing the ratio of neurons with an increase in response above threshold (Figure 2G and N). The threshold was set as the standard deviation of the population’s response to no sound and no laser power. An activity sparseness value of 0% indicates that all neurons in a population are active, and a value of 100% indicates that none of the neurons are active.

### Separation angle and Vector length

To quantify whether mean population vectors were collinear in the neuronal space, we calculated the separation angle between mean population vectors adapted from (Vinje and Gallant, 2000). For each recording, we computed the mean population vectors over the fixed time window at each laser power from 0dB to each non-zero sound pressure level (Figure 3A). We then computed the angle between each pair of mean population vectors at a given laser power (Figure 3B), and represented the mean ± s.e.m (Figure 3D-E and G-F) or the mean difference in separation angle from a given laser power to no laser power (Figure 3F and I) across recordings for each sound pair. To quantify whether mean population responses were close in the neuronal space, we computed the vector length between mean population vectors (Figure 3C). For each recording, we computed the mean population vectors over the fixed time window at each laser power between all pairs of sound pressure levels (Figure 3A). We then computed the Euclidian norm of each mean population vector at a given laser power (Figure 3C), and represented the mean ± s.e.m (Figure 3J-K and M-N) or the mean difference in vector length from a given laser power to no laser power (Figure 3L and O) across recordings for each sound pair.

**FIGURE 3:**
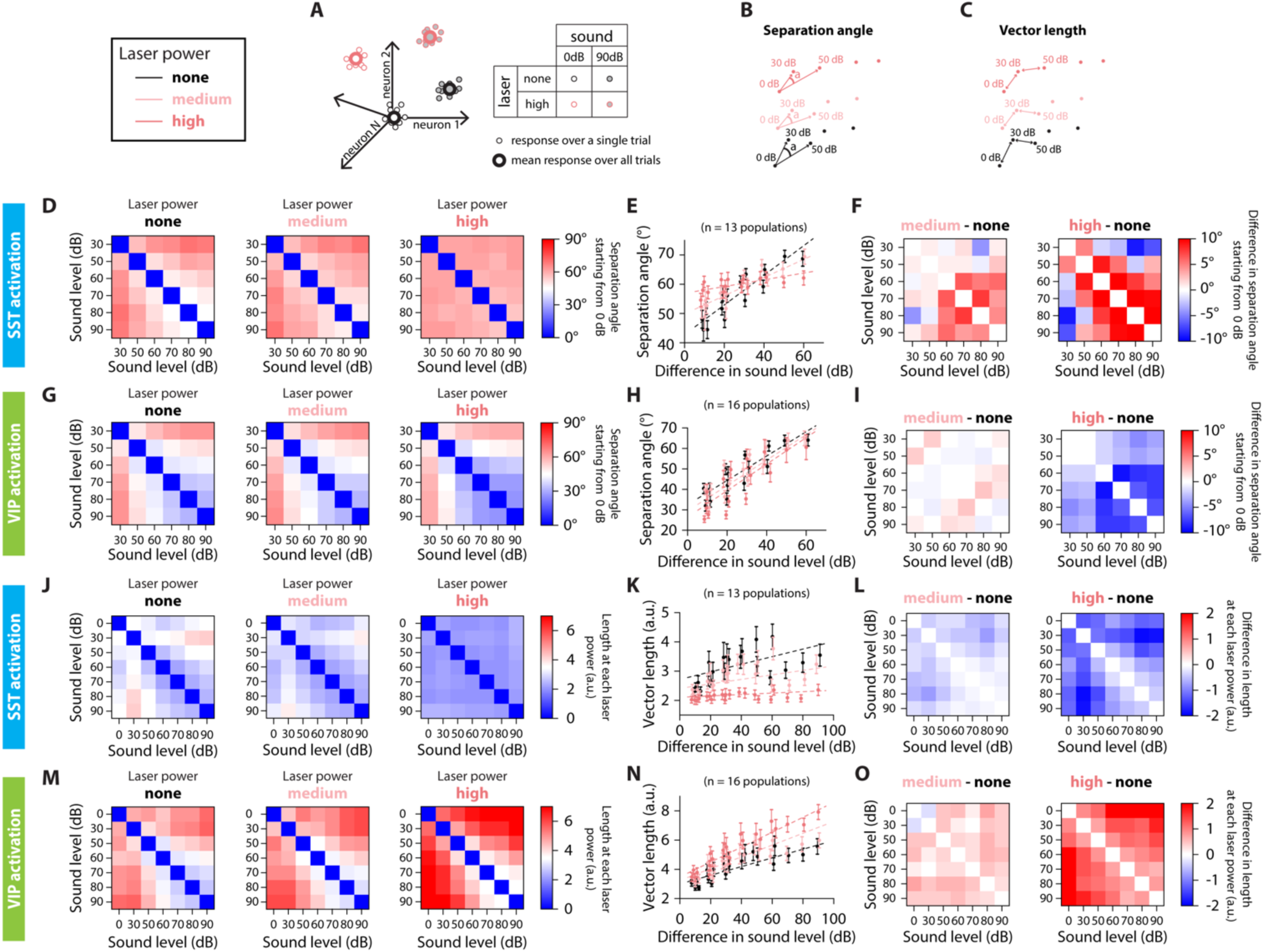
SEPARATION ANGLE AND VECTOR LENGTH ARE DIFFERENTIALLY CONTROLLED BY SST AND VIP NEURONAL ACTIVATION. A. Schematic representation of the population’s response to sound and laser stimulation, for single trials (small dots) and the average response over all trials (large dots). Sound pressure level is represented by the shading of the dot, laser power by the outline of the dot. We simplify the representation of the population’s response to two dimensions, keeping only the color code for the laser power. B. Low-dimensional schematic of the separation angle between mean population vectors to each sound pressure level at a given laser power, starting from 0dB at each laser power. C. Low-dimensional schematic of the vector length of mean population vectors between sound pressure levels at a given laser power. D. Confusion matrix of the separation angle between pairs of sound stimuli for no (left), medium (middle) and high (right) laser power of SST neuronal activation. E. Separation angle between pairs of mean population vectors to different sound pressure levels as a function of the difference in sound pressure level for no, medium and high laser powers of SST neuronal activation (circles). The dotted lines are the result from the GLME fit at the different laser powers of SST neuronal activation. F. Confusion matrix of the difference in separation angle between pairs of sound stimuli from medium (left) or high (right) laser power to no laser power of SST neuronal activation. G. Confusion matrix of the separation angle between pairs of sound stimuli for no (left), medium (middle) and high (right) laser power of VIP neuronal activation. H. Separation angle between pairs of mean population vectors to different sound pressure levels as a function of the difference in sound level for no, medium and high laser powers of VIP neuronal activation (circles). The dotted lines are the result from the GLME fit at the different laser powers of VIP neuronal activation. I. Confusion matrix of the difference in separation angle between pairs of sound stimuli from medium (left) or high (right) laser power to no laser power of VIP neuronal activation. J. Confusion matrix of the vector length between pairs of sound stimuli for no (left), medium (middle) and high (right) laser power of SST neuronal activation. K. Length of the mean population vector between pairs of sound pressure levels as a function of the difference in sound pressure level for no, medium and high laser powers of SST neuronal activation (circles). The dotted lines are the result from the GLME fit at the different laser powers of SST neuronal activation. L. Confusion matrix of the difference in vector length between pairs of sound stimuli from medium (left) or high (right) laser power to no laser power of SST neuronal activation. M. Confusion matrix of the vector length between pairs of sound stimuli for no (left), medium (middle) and high (right) laser power of VIP neuronal activation. N. Length of the mean population vector between pairs of sound pressure levels as a function of the difference in sound pressure level for no, medium and high laser powers of VIP neuronal activation (circles). The dotted lines are the result from the GLME fit at the different laser powers of VIP neuronal activation. O. Confusion matrix of the difference in vector length between pairs of sound stimuli from medium (left) or high (right) laser power to no laser power of VIP neuronal activation. For all panels, black, pink and red colors correspond to no laser power (0 mW/mm2), medium laser power (∼0.3 mW/mm2) and high laser power (∼3.5 mW/mm2), respectively (see Methods for power calibration) and the mean population vectors were computed over the fixed time window.

### Fitting of response-level curves

Mean response curves and the standard error of the mean (s.e.m) for every neuron were determined by averaging over the fixed-time window its responses to all 10 trials of each sound pressure level. Thus, for a given cell we constructed three response curves, one for every light condition.

To characterize responses as monotonic or nonmonotonic, we first normalized the response curves such that abs(max(response)) ≤ 1 and computed the monotonicity index (MI). This metric refers to the relative responses at higher stimulus levels (Watkins and Barbour, 2011) and was calculated from the mean curve as

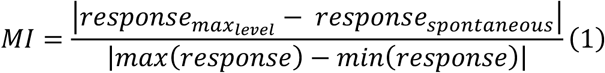

where *response_max_level__* is the response to 90 dB which is the highest level of sound presented and *response_spontaneous_* is the spontaneous response measured at 0 dB. A response curve was classified as nonmonotonic if its MI was less than 0.3 and monotonic if it was greater than 0.7. We refrain from a hard cutoff at 0.5 since preliminary analysis of the response curves indicated that due to stochasticity both monotonic and nonmonotonic curves may have MI values between 0.3 and 0.7. Furthermore, note that a given cell could change its monotonicity in the presence of optogenetic stimulation.

After determining the monotonicity of the neuronal response, we fitted the monotonic and nonmonotonic curves with a 4-parameter sigmoid function and a 4-parameter Gaussian function, respectively. The sigmoid function is given by the equation,

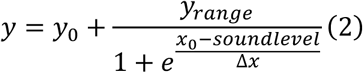

while the Gaussian function can be written as,

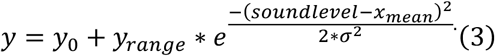

*y* refers to the response curve, *y*_0_ is the offset response, *y_range_* is its range in amplitude, *x*_0_ is the x value of the sigmoid midpoint and Δ*x* denotes the width of the sigmoid. In the Gaussian, the parameter *y*_0_ is the offset response. The amplitude, mean and standard deviation of the Gaussian are denoted by *y_range_*, *x_mean_* and σ, respectively, and have their regular interpretations. During the fitting procedure, we minimize (1 – McFadden pseudo R-squared) using the Powell optimizer (scipy.minimize.optimize in Python). The formula for this error value is,

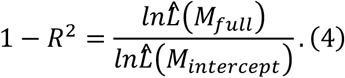

Assuming *L^*(*M_full_*) is gaussian with the experimentally computed response average as its mean value and the response s.e.m as its standard deviation, we can rewrite the formula as,

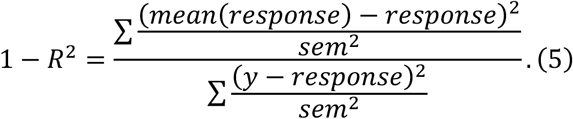

We chose the McFadden R-squared since it allows us to account for different values of s.e.m at the different intensities. The regular R-squared equation constrains the s.e.m values to be equal at all intensities. A cell’s response curve is considered well fit by its respective function if the Mcfadden *R^2^* is greater than 0.8. Due to the nonlinear nature of our optimization we chose 16 random starting points for the optimizer and cells fitted using 2 or more of the starting conditions were characterized using the fitting curve which had the highest *R*^2^ value. For neurons whose mean response curve MI lay between 0.3 and 0.7 we follow a similar procedure but fit the curve with both the sigmoid and Gaussian functions to find the better fitting function. Furthermore, we constrain the mean of all our Gaussian fits to lie between 10 and 80 dB. Response curves with means less than 10 dB or greater than 80 dB were recharacterized using the sigmoid function since only one sound pressure level data point (0 dB or 90 dB) is insufficient to adequately distinguish if the cell is monotonic or nonmonotonic. Roughly ∼5% of the total response curves (combined across all three light conditions) were refitted in this manner. Lastly, we tested if the fitting curve was overfitted to the empirical sound intensities by calculating a new variable – interpolated error. The interpolated sigmoid/Gaussian curve is constructed by interpolating the fitted curve at the intermediate sound pressure levels – (15, 40, 55, 65, 75, 85) dB. The interpolated error is the regular R-squared value evaluated using the equation,

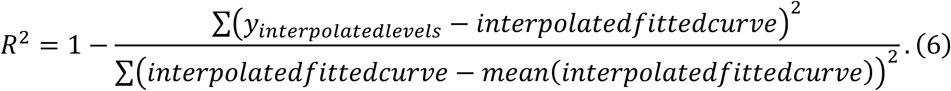

*y_interpolatedlevels_* refers to the Gaussian/sigmoid equations (2, 3) computed at the sound pressure levels 0, 15, 30, 40, 50, 55, 60, 65, 70, 75, 80, 85, and 90 dB. When computing statistics on different parameters we remove neurons with interpolated error less than 0.25 for both the Sigmoid and Gaussian fits. This threshold allows us to select at least 90% of the fitted curves.

### Decoding sound pressure level using an SVM Decoder

We linearly decoded the 7 different sound pressure levels at each opto-stimulated condition using an SVM decoder with a linear kernel. Specifically, we decoded each individual pressure level versus the remaining six. Input to the SVM consisted of the fixed time-window responses of all neurons in the population.

To individually decode each of the 7 amplitudes, for every given experimental dataset we projected the average responses of all neurons derived using a fixed window onto a lower-dimensional space using PCA. The lower-dimensional space had (n) dimensions such that these dimensions accounted for 70% of the variance in the dataset.

Next, because at a given opto-stimulated condition we had 10 trials per sound pressure level, the input data to the SVM decoder was unbalanced as 10:60. We balanced the dataset by oversampling the sound pressure level of interest, i.e., if we were decoding the 0dB stimulus from the rest, we oversampled to construct 60 trials for the 0dB stimulus. Specifically, we constructed these 60 trials by fitting the 10 experimentally obtained 0dB trials using a Gaussian kernel and sampling the corresponding distribution. The resulting oversampled dataset comprising 120 trials total was input into the two-class SVM decoder which was trained using 10-fold cross-validation. Because we were not tuning the hyperparameters of the SVM, we did not have separate validation and test sets, rather we computed the model’s accuracy directly using the validation set which was chosen randomly for each of the 10 iterations. Figure panels 2F and 2M illustrate the decoding accuracies for all 3 conditions of opto-stimulation of SST and VIP interneurons.

### Statistics

All responses are plotted as mean ± s.e.m (standard error of the mean) *with the number of measurements above the figure.* We tested significance with a Generalized Linear Mixed Effects (GLME) Model with the matlab function fitglme, using laser power, sound pressure level and the interaction term between laser power and sound pressure level as fixed-effect terms and cell identity or session number as grouping variables. All results from the statistical analyses are reported in Table 2.

**Table 2.**
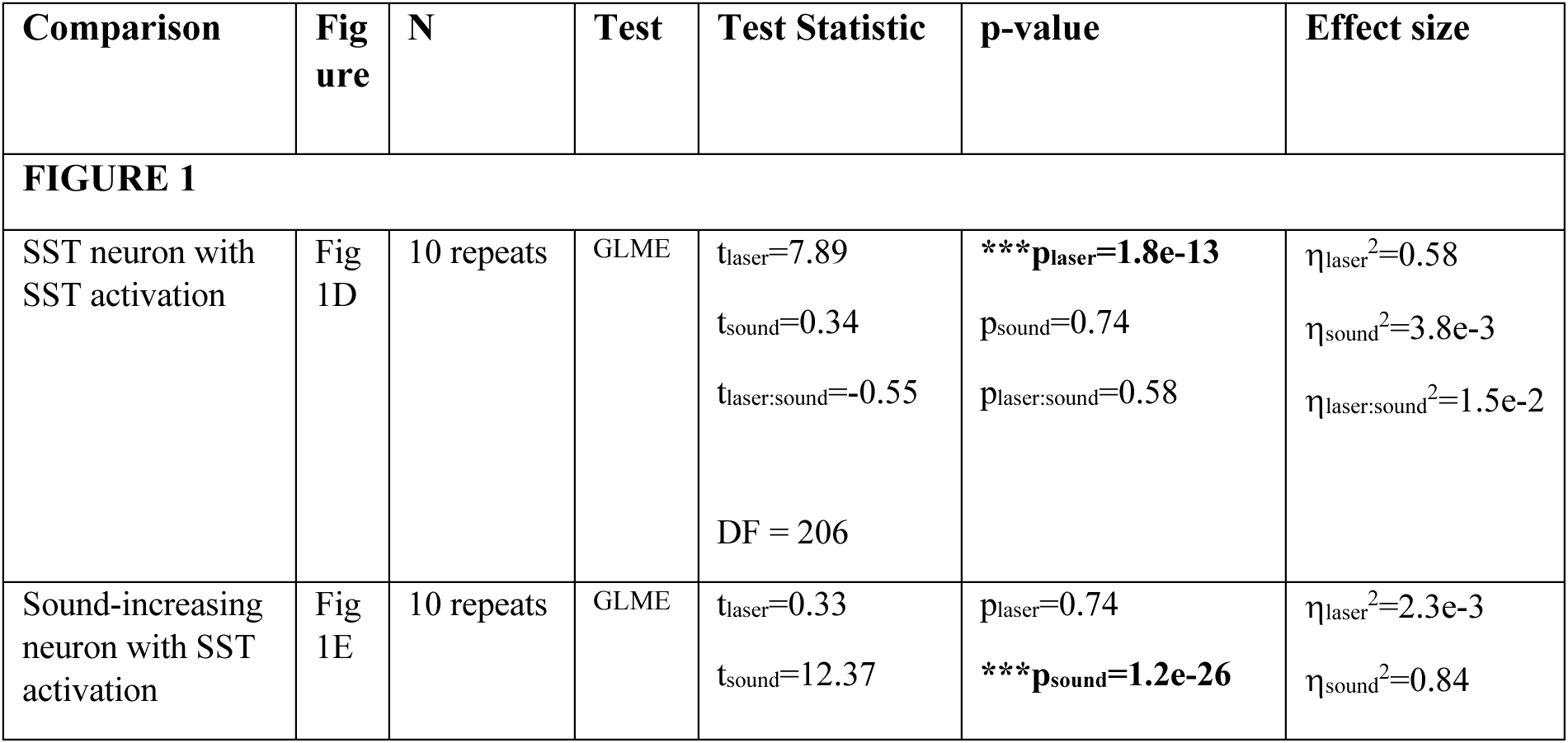
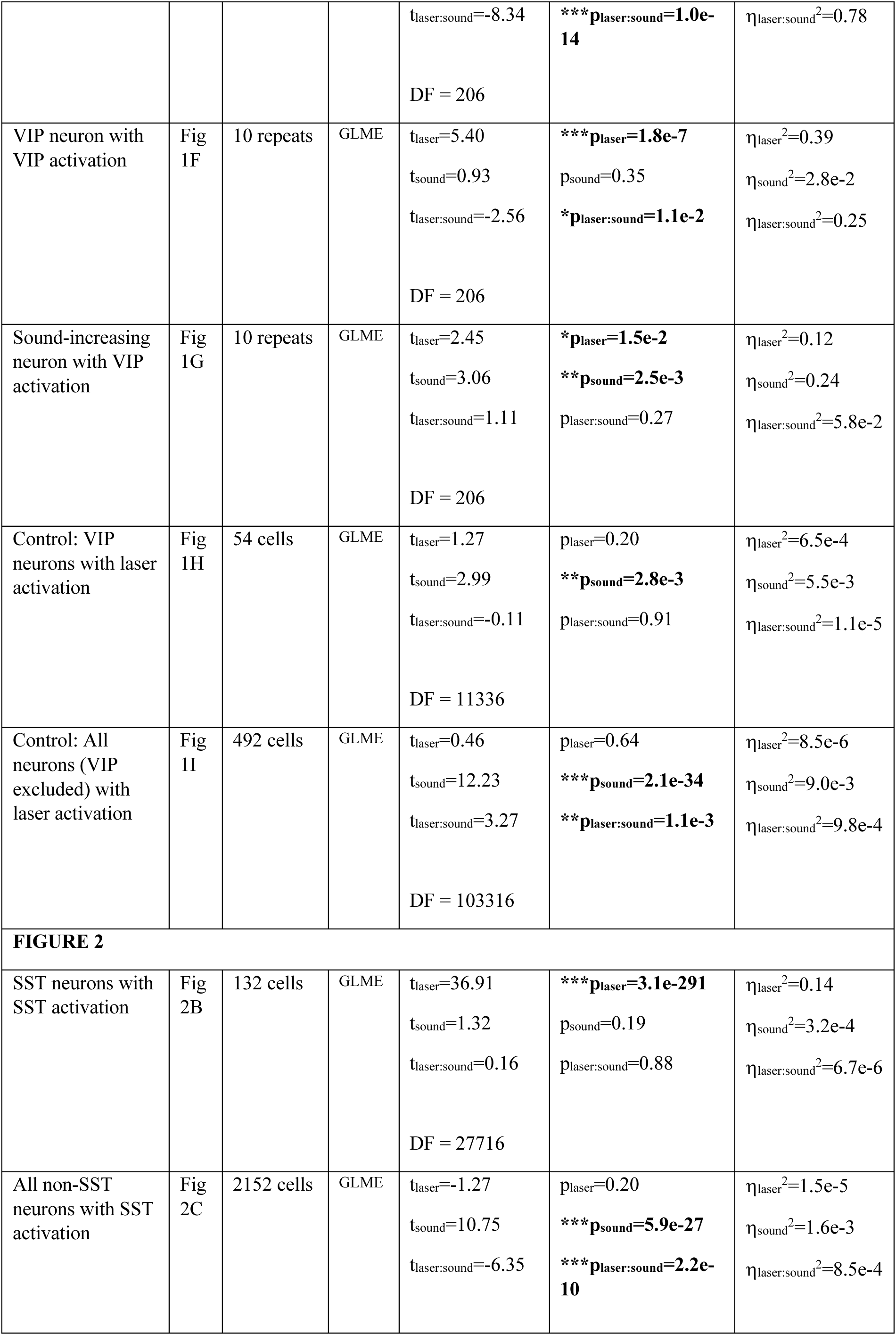
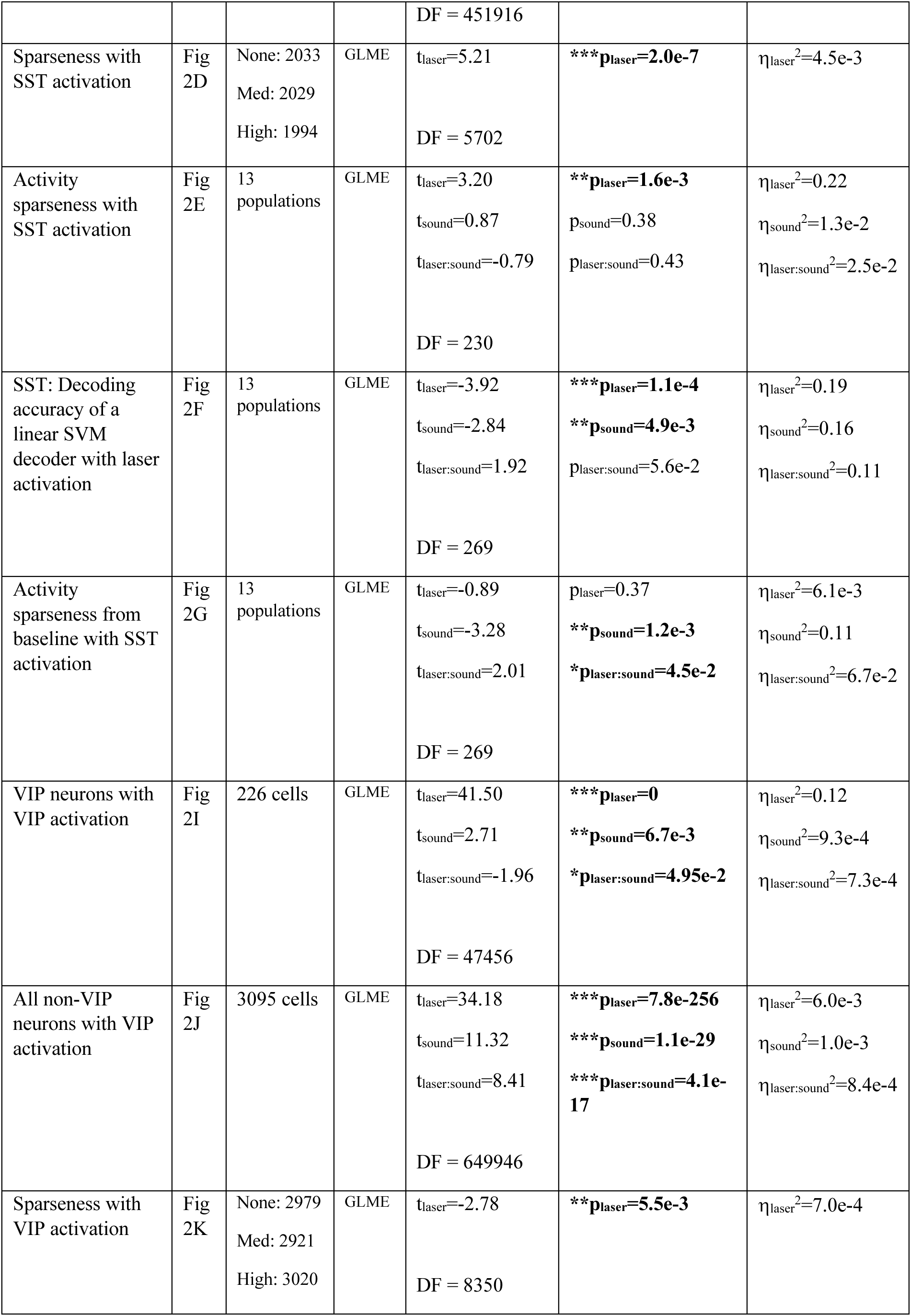
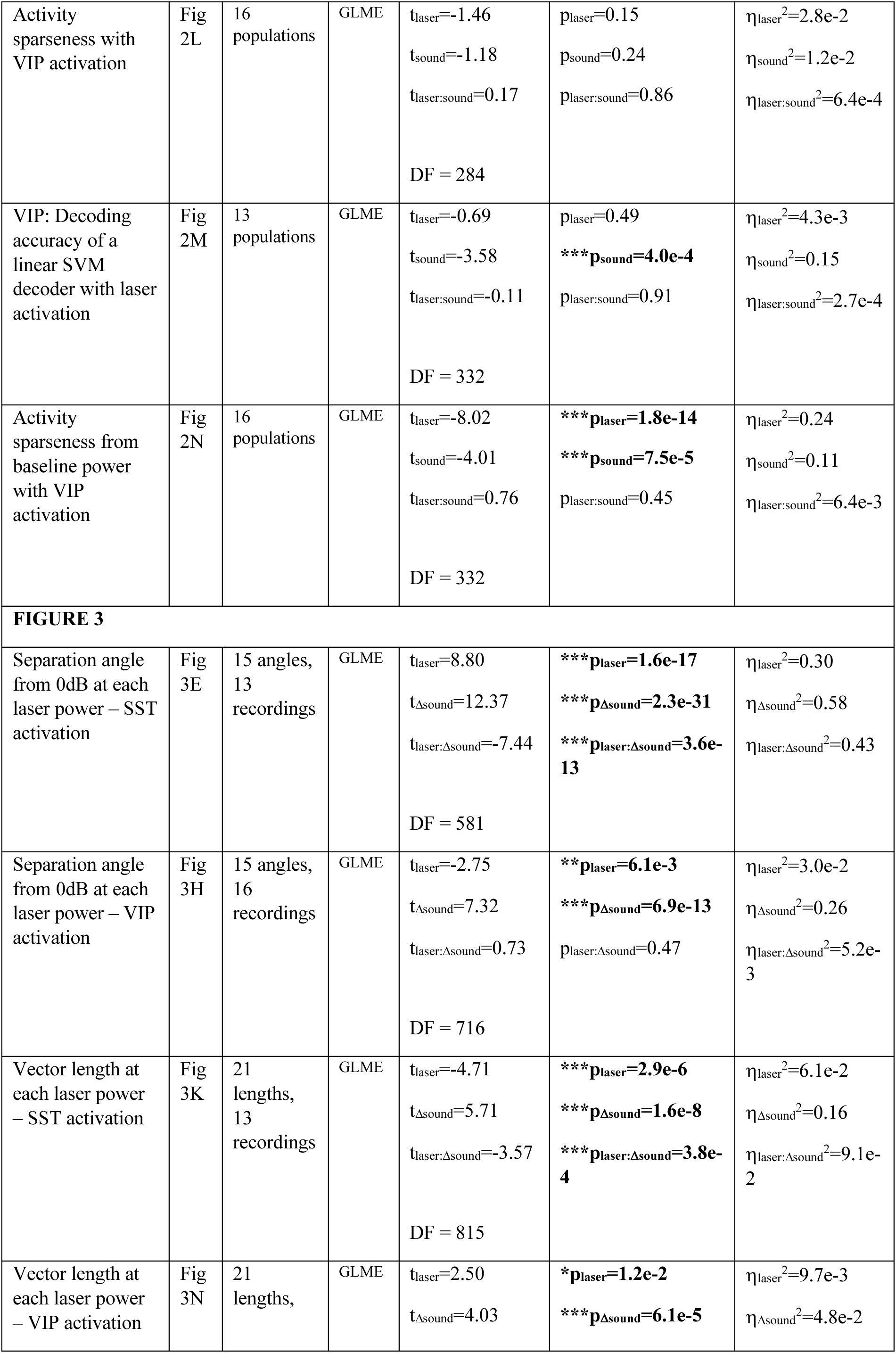
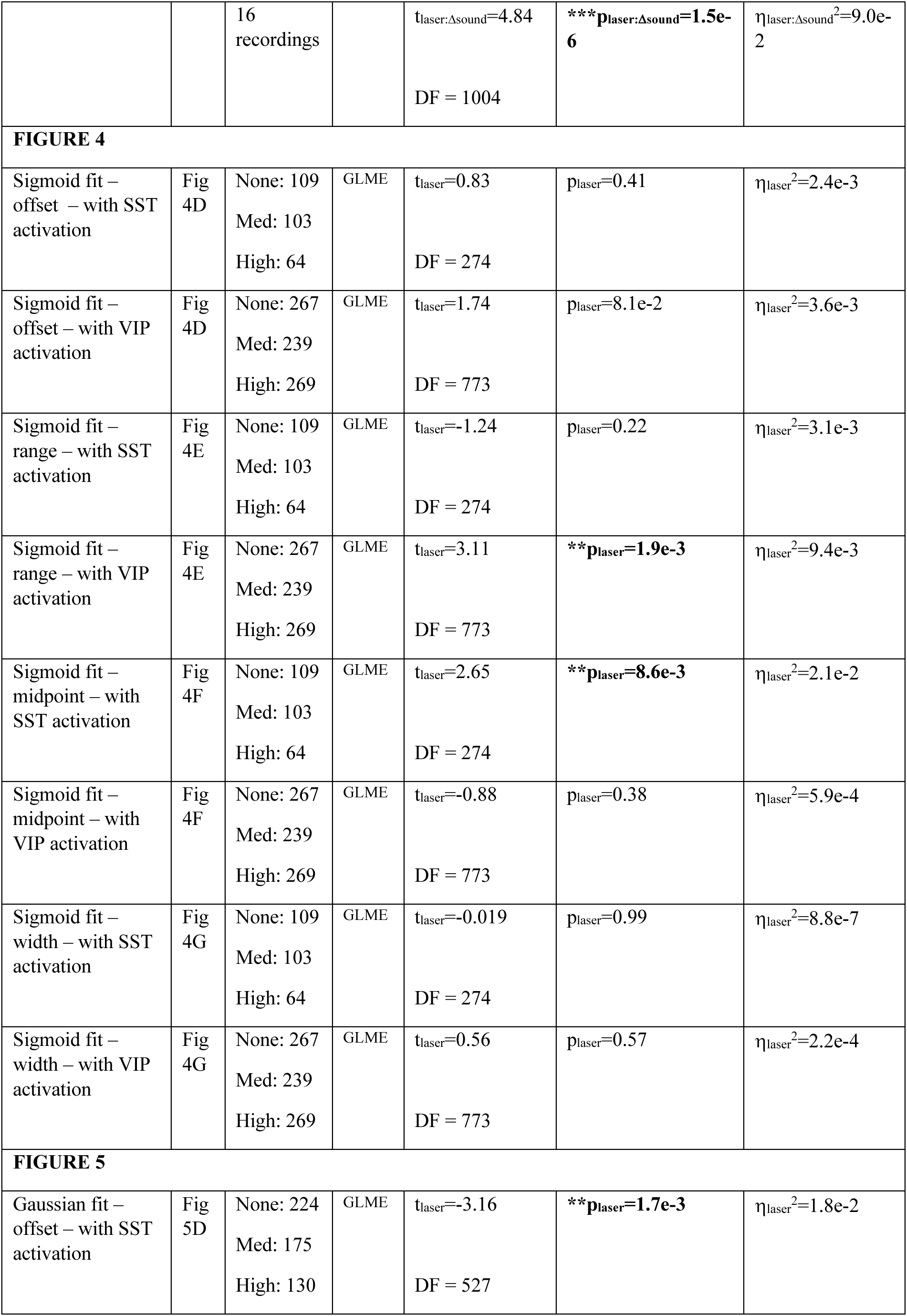
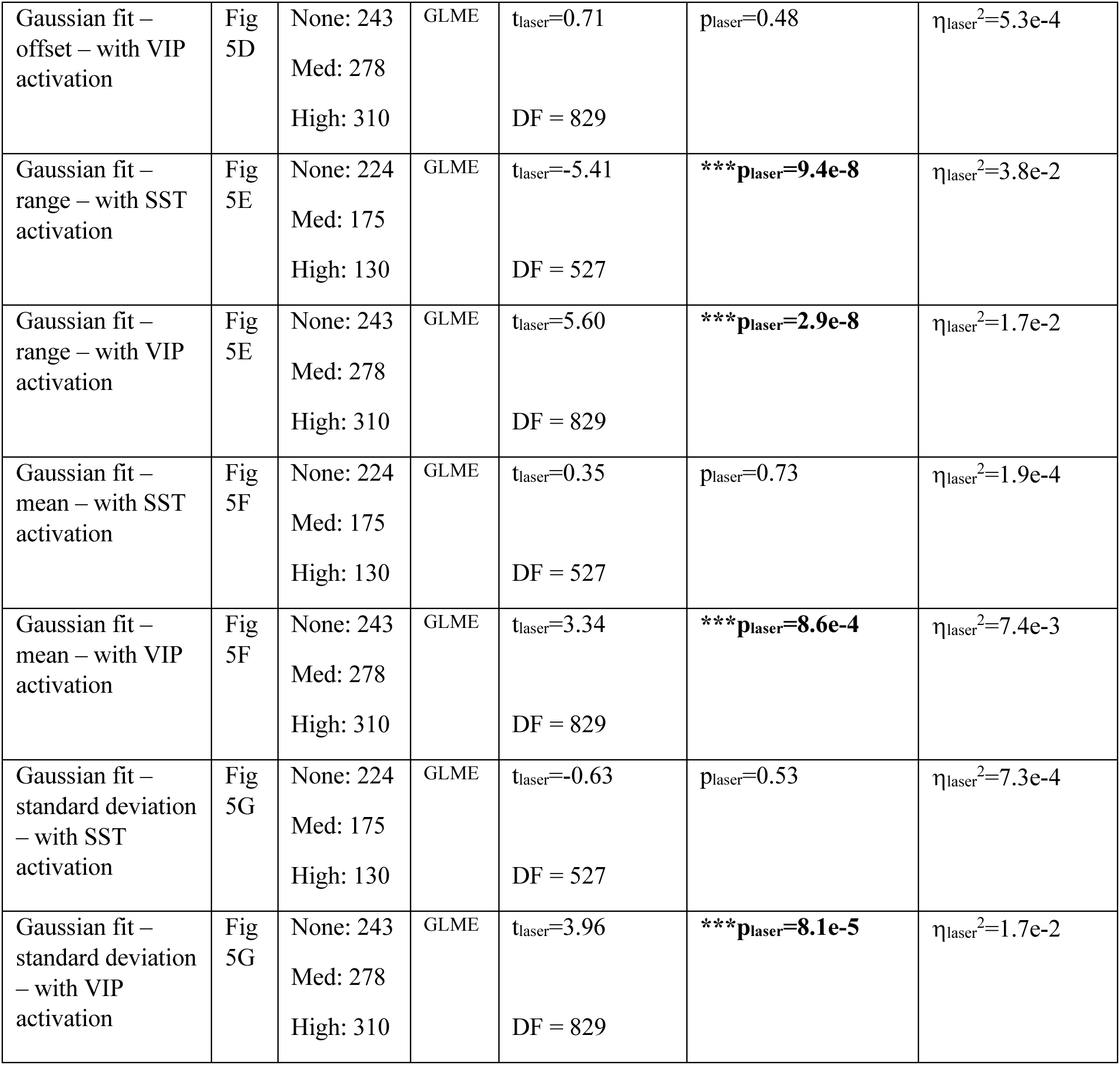
Statistics Table. We used a Generalized Linear Mixed-Effects (GLME) model and Wilcoxon signed-rank tests to compute the statistics for the data. For Figure 1, Figure 2B,C,E,G,I,J,L,N; the data (‘`table`‘) had four columns: cell, sound level, laser power, output. The formula used was (Matlab): `glme=fitglme(table,’output ∼ sound + laser + sound*laser + (1|cell)’);` For Figure 3D,K, Figure 4 and Figure 5, the data (‘`table`‘) had three columns: cell, laser power, output. The formula used was (Matlab): `glme=fitglme(table,’output ∼ laser + (1|cell)’);` For Figure 3E,H,K,N, the data (‘`table`‘) had four columns: cell, sound level difference, laser power, output. The formula used was (Matlab): `glme=fitglme(table,’output ∼ sounddiff + laser + sounddiff*laser + (1|cell)’);` For Figure 2F,M, we compared each sound amplitude across different light conditions using Wilcoxon tests.

| Comparison | Figure | N | Test | Test Statistic | p-value | Effect size |
| --- | --- | --- | --- | --- | --- | --- |
| **FIGURE 1** |  |  |  |  |  |  |
| SST neuron with SST activation | Fig 1D | 10 repeats | GLME | $t_{\text{laser}}=7.89$<br>$t_{\text{sound}}=0.34$<br>$t_{\text{laser:sound}}=-0.55$<br>DF = 206 | **\*\*\*$p_{\text{laser}}=1.8\text{e-}13$**<br>$p_{\text{sound}}=0.74$<br>$p_{\text{laser:sound}}=0.58$ | $\eta_{\text{laser}}^2=0.58$<br>$\eta_{\text{sound}}^2=3.8\text{e-}3$<br>$\eta_{\text{laser:sound}}^2=1.5\text{e-}2$ |
| Sound-increasing neuron with SST activation | Fig 1E | 10 repeats | GLME | $t_{\text{laser}}=0.33$<br>$t_{\text{sound}}=12.37$ | $p_{\text{laser}}=0.74$<br>**\*\*\*$p_{\text{sound}}=1.2\text{e-}26$** | $\eta_{\text{laser}}^2=2.3\text{e-}3$<br>$\eta_{\text{sound}}^2=0.84$ |
| | | | | $t_{\text{laser:sound}}=-8.34$<br>DF = 206 | **\*\*\*$p_{\text{laser:sound}}=1.0\text{e-}14$** | $\eta_{\text{laser:sound}}^2=0.78$ |
| VIP neuron with VIP activation | Fig 1F | 10 repeats | GLME | $t_{\text{laser}}=5.40$<br>$t_{\text{sound}}=0.93$<br>$t_{\text{laser:sound}}=-2.56$<br>DF = 206 | **\*\*\*$p_{\text{laser}}=1.8\text{e-}7$**<br>$p_{\text{sound}}=0.35$<br>**\*$p_{\text{laser:sound}}=1.1\text{e-}2$** | $\eta_{\text{laser}}^2=0.39$<br>$\eta_{\text{sound}}^2=2.8\text{e-}2$<br>$\eta_{\text{laser:sound}}^2=0.25$ |
| Sound-increasing neuron with VIP activation | Fig 1G | 10 repeats | GLME | $t_{\text{laser}}=2.45$<br>$t_{\text{sound}}=3.06$<br>$t_{\text{laser:sound}}=1.11$<br>DF = 206 | **\*$p_{\text{laser}}=1.5\text{e-}2$**<br>**\*\*$p_{\text{sound}}=2.5\text{e-}3$**<br>$p_{\text{laser:sound}}=0.27$ | $\eta_{\text{laser}}^2=0.12$<br>$\eta_{\text{sound}}^2=0.24$<br>$\eta_{\text{laser:sound}}^2=5.8\text{e-}2$ |
| Control: VIP neurons with laser activation | Fig 1H | 54 cells | GLME | $t_{\text{laser}}=1.27$<br>$t_{\text{sound}}=2.99$<br>$t_{\text{laser:sound}}=-0.11$<br>DF = 11336 | $p_{\text{laser}}=0.20$<br>**\*\*$p_{\text{sound}}=2.8\text{e-}3$**<br>$p_{\text{laser:sound}}=0.91$ | $\eta_{\text{laser}}^2=6.5\text{e-}4$<br>$\eta_{\text{sound}}^2=5.5\text{e-}3$<br>$\eta_{\text{laser:sound}}^2=1.1\text{e-}5$ |
| Control: All neurons (VIP excluded) with laser activation | Fig 1I | 492 cells | GLME | $t_{\text{laser}}=0.46$<br>$t_{\text{sound}}=12.23$<br>$t_{\text{laser:sound}}=3.27$<br>DF = 103316 | $p_{\text{laser}}=0.64$<br>**\*\*\*$p_{\text{sound}}=2.1\text{e-}34$**<br>**\*\*$p_{\text{laser:sound}}=1.1\text{e-}3$** | $\eta_{\text{laser}}^2=8.5\text{e-}6$<br>$\eta_{\text{sound}}^2=9.0\text{e-}3$<br>$\eta_{\text{laser:sound}}^2=9.8\text{e-}4$ |
| **FIGURE 2** |  |  |  |  |  |  |
| SST neurons with SST activation | Fig 2B | 132 cells | GLME | $t_{\text{laser}}=36.91$<br>$t_{\text{sound}}=1.32$<br>$t_{\text{laser:sound}}=0.16$<br>DF = 27716 | **\*\*\*$p_{\text{laser}}=3.1\text{e-}291$**<br>$p_{\text{sound}}=0.19$<br>$p_{\text{laser:sound}}=0.88$ | $\eta_{\text{laser}}^2=0.14$<br>$\eta_{\text{sound}}^2=3.2\text{e-}4$<br>$\eta_{\text{laser:sound}}^2=6.7\text{e-}6$ |
| All non-SST neurons with SST activation | Fig 2C | 2152 cells | GLME | $t_{\text{laser}}=-1.27$<br>$t_{\text{sound}}=10.75$<br>$t_{\text{laser:sound}}=-6.35$ | $p_{\text{laser}}=0.20$<br>**\*\*\*$p_{\text{sound}}=5.9\text{e-}27$**<br>**\*\*\*$p_{\text{laser:sound}}=2.2\text{e-}10$** | $\eta_{\text{laser}}^2=1.5\text{e-}5$<br>$\eta_{\text{sound}}^2=1.6\text{e-}3$<br>$\eta_{\text{laser:sound}}^2=8.5\text{e-}4$ |
|  |  |  |  | DF = 451916 |  |  |
| Sparseness with SST activation | Fig 2D | None: 2033<br>Med: 2029<br>High: 1994 | GLME | $t_{\text{laser}}=5.21$<br>DF = 5702 | **\*\*\*$p_{\text{laser}}=2.0\text{e-}7$** | $\eta_{\text{laser}}^2=4.5\text{e-}3$ |
| Activity sparseness with SST activation | Fig 2E | 13 populations | GLME | $t_{\text{laser}}=3.20$<br>$t_{\text{sound}}=0.87$<br>$t_{\text{laser:sound}}=-0.79$<br>DF = 230 | **\*\*$p_{\text{laser}}=1.6\text{e-}3$**<br>$p_{\text{sound}}=0.38$<br>$p_{\text{laser:sound}}=0.43$ | $\eta_{\text{laser}}^2=0.22$<br>$\eta_{\text{sound}}^2=1.3\text{e-}2$<br>$\eta_{\text{laser:sound}}^2=2.5\text{e-}2$ |
| SST: Decoding accuracy of a linear SVM decoder with laser activation | Fig 2F | 13 populations | GLME | $t_{\text{laser}}=-3.92$<br>$t_{\text{sound}}=-2.84$<br>$t_{\text{laser:sound}}=1.92$<br>DF = 269 | **\*\*\*$p_{\text{laser}}=1.1\text{e-}4$**<br>**\*\*$p_{\text{sound}}=4.9\text{e-}3$**<br>$p_{\text{laser:sound}}=5.6\text{e-}2$ | $\eta_{\text{laser}}^2=0.19$<br>$\eta_{\text{sound}}^2=0.16$<br>$\eta_{\text{laser:sound}}^2=0.11$ |
| Activity sparseness from baseline with SST activation | Fig 2G | 13 populations | GLME | $t_{\text{laser}}=-0.89$<br>$t_{\text{sound}}=-3.28$<br>$t_{\text{laser:sound}}=2.01$<br>DF = 269 | $p_{\text{laser}}=0.37$<br>**\*\*$p_{\text{sound}}=1.2\text{e-}3$**<br>**\*$p_{\text{laser:sound}}=4.5\text{e-}2$** | $\eta_{\text{laser}}^2=6.1\text{e-}3$<br>$\eta_{\text{sound}}^2=0.11$<br>$\eta_{\text{laser:sound}}^2=6.7\text{e-}2$ |
| VIP neurons with VIP activation | Fig 2I | 226 cells | GLME | $t_{\text{laser}}=41.50$<br>$t_{\text{sound}}=2.71$<br>$t_{\text{laser:sound}}=-1.96$<br>DF = 47456 | **\*\*\*$p_{\text{laser}}=0$**<br>**\*\*$p_{\text{sound}}=6.7\text{e-}3$**<br>**\*$p_{\text{laser:sound}}=4.95\text{e-}2$** | $\eta_{\text{laser}}^2=0.12$<br>$\eta_{\text{sound}}^2=9.3\text{e-}4$<br>$\eta_{\text{laser:sound}}^2=7.3\text{e-}4$ |
| All non-VIP neurons with VIP activation | Fig 2J | 3095 cells | GLME | $t_{\text{laser}}=34.18$<br>$t_{\text{sound}}=11.32$<br>$t_{\text{laser:sound}}=8.41$<br>DF = 649946 | **\*\*\*$p_{\text{laser}}=7.8\text{e-}256$**<br>**\*\*\*$p_{\text{sound}}=1.1\text{e-}29$**<br>**\*\*\*$p_{\text{laser:sound}}=4.1\text{e-}17$** | $\eta_{\text{laser}}^2=6.0\text{e-}3$<br>$\eta_{\text{sound}}^2=1.0\text{e-}3$<br>$\eta_{\text{laser:sound}}^2=8.4\text{e-}4$ |
| Sparseness with VIP activation | Fig 2K | None: 2979<br>Med: 2921<br>High: 3020 | GLME | $t_{\text{laser}}=-2.78$<br>DF = 8350 | **\*\*$p_{\text{laser}}=5.5\text{e-}3$** | $\eta_{\text{laser}}^2=7.0\text{e-}4$ |
| Activity sparseness with VIP activation | Fig 2L | 16 populations | GLME | $t_{\text{laser}}=-1.46$<br>$t_{\text{sound}}=-1.18$<br>$t_{\text{laser:sound}}=0.17$<br>DF = 284 | $p_{\text{laser}}=0.15$<br>$p_{\text{sound}}=0.24$<br>$p_{\text{laser:sound}}=0.86$ | $\eta_{\text{laser}}^2=2.8\text{e-}2$<br>$\eta_{\text{sound}}^2=1.2\text{e-}2$<br>$\eta_{\text{laser:sound}}^2=6.4\text{e-}4$ |
| VIP: Decoding accuracy of a linear SVM decoder with laser activation | Fig 2M | 13 populations | GLME | $t_{\text{laser}}=-0.69$<br>$t_{\text{sound}}=-3.58$<br>$t_{\text{laser:sound}}=-0.11$<br>DF = 332 | $p_{\text{laser}}=0.49$<br>**\*\*\*$p_{\text{sound}}=4.0\text{e-}4$**<br>$p_{\text{laser:sound}}=0.91$ | $\eta_{\text{laser}}^2=4.3\text{e-}3$<br>$\eta_{\text{sound}}^2=0.15$<br>$\eta_{\text{laser:sound}}^2=2.7\text{e-}4$ |
| Activity sparseness from baseline power with VIP activation | Fig 2N | 16 populations | GLME | $t_{\text{laser}}=-8.02$<br>$t_{\text{sound}}=-4.01$<br>$t_{\text{laser:sound}}=0.76$<br>DF = 332 | **\*\*\*$p_{\text{laser}}=1.8\text{e-}14$**<br>**\*\*\*$p_{\text{sound}}=7.5\text{e-}5$**<br>$p_{\text{laser:sound}}=0.45$ | $\eta_{\text{laser}}^2=0.24$<br>$\eta_{\text{sound}}^2=0.11$<br>$\eta_{\text{laser:sound}}^2=6.4\text{e-}3$ |
| **FIGURE 3** |  |  |  |  |  |  |
| Separation angle from 0dB at each laser power – SST activation | Fig 3E | 15 angles, 13 recordings | GLME | $t_{\text{laser}}=8.80$<br>$t_{\Delta\text{sound}}=12.37$<br>$t_{\text{laser}:\Delta\text{sound}}=-7.44$<br>DF = 581 | **\*\*\*$p_{\text{laser}}=1.6\text{e-}17$**<br>**\*\*\*$p_{\Delta\text{sound}}=2.3\text{e-}31$**<br>**\*\*\*$p_{\text{laser}:\Delta\text{sound}}=3.6\text{e-}13$** | $\eta_{\text{laser}}^2=0.30$<br>$\eta_{\Delta\text{sound}}^2=0.58$<br>$\eta_{\text{laser}:\Delta\text{sound}}^2=0.43$ |
| Separation angle from 0dB at each laser power – VIP activation | Fig 3H | 15 angles, 16 recordings | GLME | $t_{\text{laser}}=-2.75$<br>$t_{\Delta\text{sound}}=7.32$<br>$t_{\text{laser}:\Delta\text{sound}}=0.73$<br>DF = 716 | **\*\*$p_{\text{laser}}=6.1\text{e-}3$**<br>**\*\*\*$p_{\Delta\text{sound}}=6.9\text{e-}13$**<br>$p_{\text{laser}:\Delta\text{sound}}=0.47$ | $\eta_{\text{laser}}^2=3.0\text{e-}2$<br>$\eta_{\Delta\text{sound}}^2=0.26$<br>$\eta_{\text{laser}:\Delta\text{sound}}^2=5.2\text{e-}3$ |
| Vector length at each laser power – SST activation | Fig 3K | 21 lengths, 13 recordings | GLME | $t_{\text{laser}}=-4.71$<br>$t_{\Delta\text{sound}}=5.71$<br>$t_{\text{laser}:\Delta\text{sound}}=-3.57$<br>DF = 815 | **\*\*\*$p_{\text{laser}}=2.9\text{e-}6$**<br>**\*\*\*$p_{\Delta\text{sound}}=1.6\text{e-}8$**<br>**\*\*\*$p_{\text{laser}:\Delta\text{sound}}=3.8\text{e-}4$** | $\eta_{\text{laser}}^2=6.1\text{e-}2$<br>$\eta_{\Delta\text{sound}}^2=0.16$<br>$\eta_{\text{laser}:\Delta\text{sound}}^2=9.1\text{e-}2$ |
| Vector length at each laser power – VIP activation | Fig 3N | 21 lengths, | GLME | $t_{\text{laser}}=2.50$<br>$t_{\Delta\text{sound}}=4.03$ | **\*$p_{\text{laser}}=1.2\text{e-}2$**<br>**\*\*\*$p_{\Delta\text{sound}}=6.1\text{e-}5$** | $\eta_{\text{laser}}^2=9.7\text{e-}3$<br>$\eta_{\Delta\text{sound}}^2=4.8\text{e-}2$ |
| | | 16 recordings | | $t_{\text{laser}:\Delta\text{sound}}=4.84$<br>DF = 1004 | **\*\*\*$p_{\text{laser}:\Delta\text{sound}}=1.5\text{e-}6$** | $\eta_{\text{laser}:\Delta\text{sound}}^2=9.0\text{e-}2$ |
| **FIGURE 4** |  |  |  |  |  |  |
| Sigmoid fit – offset – with SST activation | Fig 4D | None: 109<br>Med: 103<br>High: 64 | GLME | $t_{\text{laser}}=0.83$<br>DF = 274 | $p_{\text{laser}}=0.41$ | $\eta_{\text{laser}}^2=2.4\text{e-}3$ |
| Sigmoid fit – offset – with VIP activation | Fig 4D | None: 267<br>Med: 239<br>High: 269 | GLME | $t_{\text{laser}}=1.74$<br>DF = 773 | $p_{\text{laser}}=8.1\text{e-}2$ | $\eta_{\text{laser}}^2=3.6\text{e-}3$ |
| Sigmoid fit – range – with SST activation | Fig 4E | None: 109<br>Med: 103<br>High: 64 | GLME | $t_{\text{laser}}=-1.24$<br>DF = 274 | $p_{\text{laser}}=0.22$ | $\eta_{\text{laser}}^2=3.1\text{e-}3$ |
| Sigmoid fit – range – with VIP activation | Fig 4E | None: 267<br>Med: 239<br>High: 269 | GLME | $t_{\text{laser}}=3.11$<br>DF = 773 | **\*\*$p_{\text{laser}}=1.9\text{e-}3$** | $\eta_{\text{laser}}^2=9.4\text{e-}3$ |
| Sigmoid fit – midpoint – with SST activation | Fig 4F | None: 109<br>Med: 103<br>High: 64 | GLME | $t_{\text{laser}}=2.65$<br>DF = 274 | **\*\*$p_{\text{laser}}=8.6\text{e-}3$** | $\eta_{\text{laser}}^2=2.1\text{e-}2$ |
| Sigmoid fit – midpoint – with VIP activation | Fig 4F | None: 267<br>Med: 239<br>High: 269 | GLME | $t_{\text{laser}}=-0.88$<br>DF = 773 | $p_{\text{laser}}=0.38$ | $\eta_{\text{laser}}^2=5.9\text{e-}4$ |
| Sigmoid fit – width – with SST activation | Fig 4G | None: 109<br>Med: 103<br>High: 64 | GLME | $t_{\text{laser}}=-0.019$<br>DF = 274 | $p_{\text{laser}}=0.99$ | $\eta_{\text{laser}}^2=8.8\text{e-}7$ |
| Sigmoid fit – width – with VIP activation | Fig 4G | None: 267<br>Med: 239<br>High: 269 | GLME | $t_{\text{laser}}=0.56$<br>DF = 773 | $p_{\text{laser}}=0.57$ | $\eta_{\text{laser}}^2=2.2\text{e-}4$ |
| **FIGURE 5** |  |  |  |  |  |  |
| Gaussian fit – offset – with SST activation | Fig 5D | None: 224<br>Med: 175<br>High: 130 | GLME | $t_{\text{laser}}=-3.16$<br>DF = 527 | **\*\*$p_{\text{laser}}=1.7\text{e-}3$** | $\eta_{\text{laser}}^2=1.8\text{e-}2$ |
| Gaussian fit –<br>offset – with VIP<br>activation | Fig<br>5D | None: 243<br>Med: 278<br>High: 310 | GLME | $t_{\text{laser}}=0.71$<br>DF = 829 | $p_{\text{laser}}=0.48$ | $\eta_{\text{laser}}^2=5.3\text{e-}4$ |
| Gaussian fit –<br>range – with SST<br>activation | Fig<br>5E | None: 224<br>Med: 175<br>High: 130 | GLME | $t_{\text{laser}}=-5.41$<br>DF = 527 | ***$p_{\text{laser}}=9.4\text{e-}8$ | $\eta_{\text{laser}}^2=3.8\text{e-}2$ |
| Gaussian fit –<br>range – with VIP<br>activation | Fig<br>5E | None: 243<br>Med: 278<br>High: 310 | GLME | $t_{\text{laser}}=5.60$<br>DF = 829 | ***$p_{\text{laser}}=2.9\text{e-}8$ | $\eta_{\text{laser}}^2=1.7\text{e-}2$ |
| Gaussian fit –<br>mean – with SST<br>activation | Fig<br>5F | None: 224<br>Med: 175<br>High: 130 | GLME | $t_{\text{laser}}=0.35$<br>DF = 527 | $p_{\text{laser}}=0.73$ | $\eta_{\text{laser}}^2=1.9\text{e-}4$ |
| Gaussian fit –<br>mean – with VIP<br>activation | Fig<br>5F | None: 243<br>Med: 278<br>High: 310 | GLME | $t_{\text{laser}}=3.34$<br>DF = 829 | ***$p_{\text{laser}}=8.6\text{e-}4$ | $\eta_{\text{laser}}^2=7.4\text{e-}3$ |
| Gaussian fit –<br>standard deviation<br>– with SST<br>activation | Fig<br>5G | None: 224<br>Med: 175<br>High: 130 | GLME | $t_{\text{laser}}=-0.63$<br>DF = 527 | $p_{\text{laser}}=0.53$ | $\eta_{\text{laser}}^2=7.3\text{e-}4$ |
| Gaussian fit –<br>standard deviation<br>– with VIP<br>activation | Fig<br>5G | None: 243<br>Med: 278<br>High: 310 | GLME | $t_{\text{laser}}=3.96$<br>DF = 829 | ***$p_{\text{laser}}=8.1\text{e-}5$ | $\eta_{\text{laser}}^2=1.7\text{e-}2$ |

### Data availability

The data and code are available on the dryad depository: https://doi.org/10.5061/dryad.t1g1jwt6d (Melanie Tobin et al., n.d.).

## RESULTS

### SST and VIP neuronal activation modulate the response of sound-increasing neurons

To investigate whether and how distinct classes of inhibitory neurons in AC, SST and VIP neurons, affect sound representation at population level, we imaged Calcium activity of neurons in AC of awake, head-fixed mice presented with sounds while activating SST or VIP neurons using sub-millisecond optogenetic manipulation with Chrimson (Figure 1A) (Klapoetke et al., 2014). We monitored calcium activity by measuring fluorescence of GCaMP expressed in hundreds of neurons at a time (VIP-Cre mice: 3321 neurons over 16 recordings, SST-Cre mice: 2284 neurons over 13 recordings) and identified the cells expressing the opsin through co-expression of tdTomato (Figure 1B and 1C). This approach allowed us to quantify the transformations of sound representations within a large population of cortical neurons driven by SST or VIP neuronal activation.

We first confirmed that the optogenetic manipulations produced expected responses. SST neurons directly inhibit excitatory cells and other cells within the neuronal population, and so we expected that optogenetic manipulation would increase SST neuronal activity, but decrease the responses of other cells. By contrast, VIP neurons mostly inhibit other inhibitory neurons (Campagnola et al., 2022), and therefore we expected the VIP neuronal activation would increase both VIP neuronal activity and provide a release of inhibition to other cells in the network. In SST-Cre mice, a representative SST neuron increased activity at all sound pressure levels with laser (example neuron, \*\*\**p*_laser_=1.8e-13, GLME, Figure 1D). The change in the response of a representative non-SST neuron to SST neuronal activation was sound level-dependent, with a decrease at most sound pressure levels for the medium laser power. Sound responses were abolished with strong SST neuronal activation at the high laser power (*p*_laser_=0.74, \*\*\**p*_laser:sound_=1.0e-14, GLME, Figure 1E). In VIP-Cre mice, the response of a representative VIP neuron increased at all sound pressure levels with laser and the increase was sound-level dependent (example neuron, \*\*\**p*_laser_=1.8e-7, \**p*_laser:sound_=1.1e-2, GLME, Figure 1F). The activity of a representative non-VIP neuron increased at most sound pressure levels during activation of VIP neurons, with a larger increase at the high than medium laser power (\**p*_laser_=1.5e-2, GLME, Figure 1G). As a control, we injected mice with Flex.tdTomato instead of the opsin in VIP-Cre x Cdh23 mice (n=5), and verified that laser stimulation of AC in the absence of the opsin did not lead to significant changes in the neuronal responses of VIP neurons (*p*_laser_=0.20, GLME, Figure 1H) and non-VIP neurons (*p*_laser_=0.64, GLME, Figure 1I). These representative effects of SST or VIP neuronal activation are consistent with previous reports, suggesting that the activation method worked as expected(Natan et al., 2015; Seybold et al., 2015; Phillips and Hasenstaub, 2016; Bigelow et al., 2019).

### Differential distribution of population activity and sparseness with SST and VIP neuronal activation

We next tested whether and how SST and VIP neuronal activation differentially affects the sound pressure level response functions of neurons in AC. We characterized the response of each population to any stimulus combination over its fixed time window (Figure 2 panels A-C and H-J, see Methods).

Because SST neurons directly inhibit excitatory cells, we hypothesized that SST neuronal activation would lead to fewer neurons responding at increasing sound pressure levels. At baseline, SST neurons have on average lower responses than the non-SST neurons, with the difference in response strongest at low sound pressure levels (Figure 2B-C). During the SST neuronal activation, the average response of SST neurons increased at all sound pressure levels (n=132 SST neurons, \*\*\**p*_laser_<1e-100, GLME, Figure 2A, solid lines Figure 2B), whereas non-SST neurons exhibited a mix of decreased and increased responses (Figure 2A). At medium and high laser, the overall shape of the average response-level curve is preserved for SST neurons (plaser:sound=0.88, GLME, Figure 2B). The overall effect of SST neuronal activation on the population of non-SST neurons had a significant interaction between sound and laser amplitude (dashed lines, Figure 2C), but no significant sound-independent laser effect (solid and dashed lines, n=2152 neurons, *p*_laser_=0.20, \*\*\**p*_laser:sound_=2.2e-10, GLME, Figure 2C). The average response-level curve for non-SST neurons shifted downwards for the medium laser power, and at a high laser power, the modulation of the population’s response by sound was lost with an average response to silence equal to the average response to sounds.

To further assess the representation of sound pressure level in the neuronal population, we studied two characteristics of sparse distributed representations: (1) each neuron responds only to a few stimuli (high sparseness) and (2) only a few neurons respond to each stimulus (high activity sparseness). To measure how many stimuli a neuron responds to, we computed the sparseness of each non-SST neuron (see Methods). The sparseness of non-SST neurons with a positive sound response increased significantly from 62% to 67% upon SST neuronal activation (median, n=2033, 2029, and 1994 neurons for no, medium and high laser power, respectively; \*\*\**p*_laser_ = 2.0e-7, GLME, Figure 2D), indicating that neurons responded to fewer stimuli. To measure how many neurons responded to each stimulus, we computed the activity sparseness for each population of non-SST neurons, which is the ratio of neurons that are *not* active in response to a given stimulus. The activity sparseness compared to silence at a given laser power increased significantly upon SST neuronal activation (n=13 populations, \*\**p*_laser_=1.6e-3, GLME, Figure 2E), indicating that at each successive sound pressure level, there were fewer non-SST neurons that were active. Compared to baseline, the activity sparseness showed a significant interaction between sound and laser amplitude but no sound-independent laser effect upon SST neuronal activation (n=13 populations, *p*_laser_=0.37, *\*p*laser:sound=4.5e-2, GLME, Figure 2G). However, whereas decoding accuracy differed across different sound pressure levels, and was slightly lower with SST neuronal activation, there was no interaction between the laser and sound amplitudes (n=13 populations, \*\**p*_sound_=4.9e-3, \*\*\**p*_laser_ =1.1e-4, *p*_laser:sound_=0.056, GLME, Figure 2F). This suggests that despite the neuronal population responses becoming relatively sparser across sound pressure levels, the relative decoding accuracy was preserved across SPLs. Combined with the change in sound modulation of the population’s response, these results point to a more localist representation for sound pressure level with SST neuronal activation.

Because VIP neuron activity provides a release of SST neuron inhibition on excitatory cells, we hypothesized that VIP neuronal activation would lead to more neurons being active in response to each sound level and having stronger responses. At baseline, VIP neurons have on average lower responses than the non-VIP neurons, with the difference being smallest at 30dB (Figure 2I-J). When VIP neurons were activated, the average response of VIP neurons increased at all sound pressure levels (n=226 VIP neurons, \*\*\**p*_laser_<1e-100, GLME, Figure 2H, solid lines Figure 2I). At high laser, the sound modulation weakens in VIP neurons (*plaser:sound=4.95e-2, GLME, Figure 2I). Non-VIP neurons similarly exhibited an increase in response, both in silence and to sounds at different sound pressure levels (Figure 2H). The average response-level curve for non-VIP neurons shifted upwards at all sound pressure levels as VIP neurons were activated (solid and dashed lines, n=3095 neurons, \*\*\**p*_laser_<1e-100, GLME, Figure 2J), reflective of a more distributed stimulus representation. At all laser powers, the modulation of the population’s response by sound was maintained: the population average to sounds was higher than to silence (dashed lines, Figure 2J).

Neuronal sparseness decreased with VIP neuronal activation for all non-VIP neurons with a positive sound response, with a significant decrease from 54% to 52% upon VIP neuronal activation (median, n=2979; 2921 and 3020 neurons for no, medium and high laser power, respectively; \*\**p*_laser_=5.5e-3, GLME, Figure 2K), indicating that each neuron responded more equally to the different sound pressure levels. The activity sparseness measured from silence at a given laser power did not change upon VIP neuronal activation (n=16 populations, *p*_laser_=0.15, GLME, Figure 2L), indicating that the same number of neurons showed an increase in response at each sound pressure level from silence at a given laser power. Consistent with the overall increase in responses with VIP neuronal activation in silence, the activity sparseness measured from baseline significantly decreased (n=16 populations, \*\*\**p*_laser_=1.8e-14, GLME, Figure 2N). Importantly, this change in population activity did not affect the decoding performance (n=13 populations, *p*_sound_=4.0e-4, *p*_laser_=0.49, *p*_laser:sound_*=*0.91, GLME, Figure 2M). Combined, these results suggest that activating VIP neurons transforms population responses to a more distributed representation, while preserving the decoding accuracy.

Overall, SST neuronal activation led to weaker and sparser responses in the population, with neurons responding to fewer stimuli and each stimulus eliciting a response in fewer neurons, shifting the population responses toward a more localist stimulus representation. By contrast VIP neuronal activation leads to a global increase in the neuronal population’s response, along with each neuron responding to more stimuli, leading to a more distributed stimulus representation.

### Sound pressure level is represented more discretely or continuously in the neuronal population with SST or VIP neuronal activation, respectively

There are various ways that a neuronal network can implement a representation of a sensory feature. For example, a distributed code may rely on the magnitude of response of the population of neurons or on the relative response of each cell. To investigate this, we examined next the properties of the representation of sound pressure level upon SST and VIP neuronal activation in the neuronal space. We computed the mean population vector from 0dB at a given laser amplitude to each nonzero sound pressure level at that laser amplitude (Figure 3A), and computed the separation angle between pairs of mean population vectors at the same laser power (Figure 3B), as well as the length of mean population vectors between pairs of sounds (Figure 3C).

The separation angle (Vinje and Gallant, 2000) computes the angle between the mean population vectors to two different sound pressure levels taking 0dB at each laser power as the origin (Figure 3B). A smaller angle indicates the population vectors are more collinear, meaning similar neurons respond to the different stimuli, although perhaps with differing magnitude. A larger angle indicates that there is less overlap between the populations of neurons responding to each stimulus.

When there is no interneuron activation, the separation angle increased with the difference in sound pressure level (Figure 3E and H, black circles), indicating that there is less overlap in the groups of neurons responding to sounds with a large difference in sound pressure level than a small difference in sound pressure level. Upon SST neuronal activation, the curve flattened around 60°, meaning that there was an increase in separation angle for small differences in sound pressure level, and a decrease in separation angle for large differences in sound pressure level (dotted lines correspond to GLME estimates, \*\*\**p*_laser_=1.6e-17, \*\*\**p*_Δsound_=2.3e-31, \*\*\**p*_laser_:Δsound=3.6e-13, GLME, Figure 3D-E). On average, the difference in separation angle between sound pairs from medium or high laser to no laser was positive, at 3.7° ± 1.7° for the high laser power (mean ± s.e.m, 15 angles, Figure 3F). This indicates that the population vectors to the different tested sound pressure levels were more equally distributed in the neuronal space. There is, however, still an overlap between groups of neurons responding to different sound pressure levels, as a fully orthogonal coding of sound pressure level would lead to a 90° separation angle. In contrast, upon VIP neuronal activation, the separation angle decreased equally for all differences in sound pressure level (dotted lines correspond to GLME estimates, \*\**p*_laser_=6.1e-3, \*\*\**p*_Δsound_=6.9e-13, *p*_laser:Δsound_=0.47, GLME, Figure 3G-H). On average, the difference in separation angle between sound pairs from medium or high laser to no laser was negative, at − 4.6° ± 0.7° degrees at the high laser power (mean ± s.e.m, 15 angles, Figure 3I). This indicates that the population vectors to the different tested sound pressure levels were more collinear, with more overlap between groups of neurons responding to different sound pressure levels with VIP neuronal activation.

The vector length computes the Euclidian norm of the mean population vector between two sound pressure levels at a given laser power (Figure 3C). A small length indicates that the responses to two different stimuli are close in the neuronal space, either due to small magnitudes of response, to small differences in separation angle or both, whereas a large length indicates that there is a large difference in magnitude, in separation angle or both. Therefore, we tested whether SST and VIP neuronal activation differentially affected the vector length.

Upon SST neuronal activation, the vector length decreased for all sound pressure level differences to about 2 a.u. at high laser power, along with a decrease in the slope of the GLME estimate by 81% at high laser power (dotted lines correspond to GLME estimates, \*\*\**p*_laser_=2.9e-6, \*\*\**p*_Δsound_=1.6e-8, \*\*\**p*_laser:Δsound_=3.8e-4, GLME, Figure 3J-K). The average change in length from medium or high laser power to no laser was negative, at −1.00 ± 0.10 a.u. for the high laser power (mean ± s.e.m, 21 lengths, Figure 3L). Upon VIP neuronal activation, the vector length increased for all sound pressure level differences, and the slope of the GLME estimate also increased by 72% at high laser power (dotted lines correspond to GLME estimates, \**p*_laser_=1.2e-2, \*\*\**p*_Δsound_=6.1e-5, \*\*\**p*_laser:Δsound_=1.5e-6, GLME, Figure 3M-N). The average change in length from medium or high laser power to no laser was positive, at 1.27 ± 0.12 a.u. for the high laser power (mean ± s.e.m, 21 lengths, Figure 3O).

Combined with the differences in the separation angle, these results point to emergence of two types of modulations of population codes: Upon SST neuronal activation, the encoding of sound pressure level resembles a localist pattern coding where the magnitude of response is less relevant than the identity of responding cells: the strength of response is reduced and similar for all sound pressure levels, but the population vector angles are more spread out in neuronal space, indicating that there is less overlap between groups of neurons responding to different sound pressure levels. Upon VIP neuronal activation, the encoding of sound pressure level resembles a rate code, which is a type of distributed representation in which the varying strength of the whole population encodes a continuously varying parameter of the stimulus. Therefore, VIP neuronal activation promotes the strength of response more than the identity of responding neurons: there is more overlap between the groups of neurons responding to different sound pressure levels, but the strength of response is increased.

### Response-level curves of sound-modulated cells exhibit a narrower response upon SST neuronal activation, and a broader response upon VIP neuronal activation

We next tested how the shifts in stimulus representation mediated by SST and VIP neurons at the scale of the neuronal population are implemented at the single-cell level by analyzing the changes with SST or VIP neuronal activation in response-level curves of single neurons responding positively to sound. Some AC neurons exhibit increased responses with increased sound pressure levels (monotonic response-level curve) while others are tuned with a peak response to a specific sound pressure level (nonmonotonic response-level curve) (Schreiner et al., 1992; Phillips et al., 1995; Wu et al., 2006). We classified cells depending on their Monotonicity Index (MI, see Methods) and fit response functions of monotonically responding cells with a Sigmoid function (Figure 4 see Methods) and those of non-monotonically responding cells with a Gaussian function (Figure 5, see Methods). We then tested how the parameters of the fits change with interneuron activation.

**FIGURE 4:**
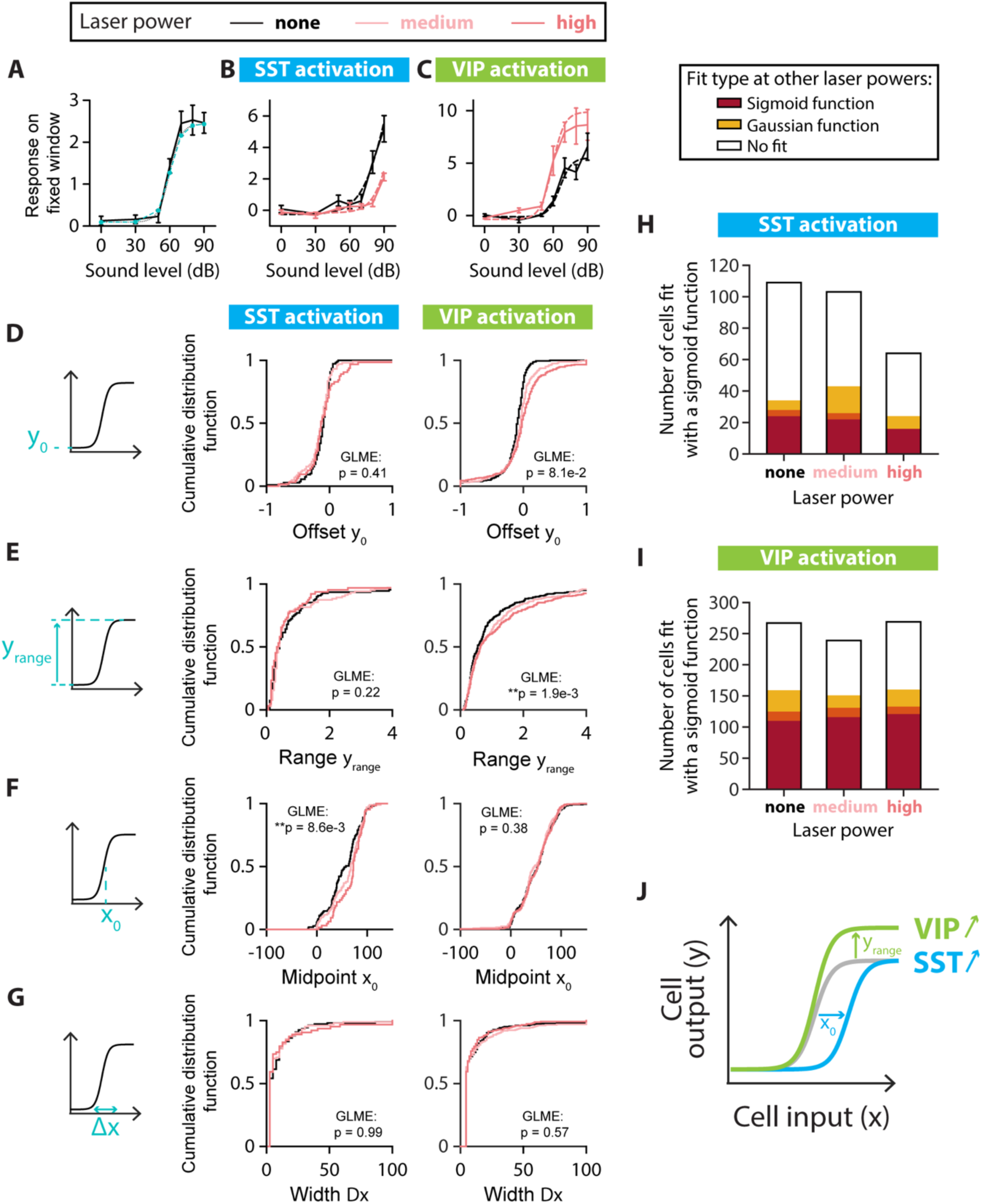
SINGLE-CELL FITS OF RESPONSE-LEVEL CURVES FOR MONOTONIC CELLS. A. Example neuron with a monotonic response-level curve (solid black line) with a sigmoid fit estimated at the probed sound amplitudes (blue dashed line with circles) and the sigmoid function with the same parameters (dotted black line). The parameters for the sigmoid fit are: Offset amplitude *y*0 = 0.1; Range *y*range = 2.4; Midpoint *x*0 = 60 dB; Width Δ*x* = 5 dB. B. Example of changes to the response-level curve of a sound-increasing monotonic neuron upon SST neuronal activation. Response-level curves (solid lines) are fit by sigmoid functions (dotted lines) at no (black lines) and high (red lines) laser powers of SST neuronal activation. The parameters for the sigmoid fit are, for no SST neuronal activation: *y*0 = -0.3; *y*range = 13.4; *x*0 = 93 dB; Δ*x* = 11 dB; and for SST neuronal activation: *y*0 = -0.2; *y*range = 8.6; *x*0 = 96 dB; Δ*x* = 8 dB. C. Example of changes to the response-level curve of a sound-increasing monotonic neuron upon VIP neuronal activation. Response-level curves (solid lines) are fit by sigmoid functions (dotted lines) at no (black lines) and high (red lines) laser powers of VIP neuronal activation. The parameters for the sigmoid fit are, for no VIP neuronal activation: *y*0 = -0.1; *y*range = 5.6; *x*0 = 66 dB; Δ*x* = 5 dB; and for VIP neuronal activation: *y*0 = -0.4; *y*range = 9.9; *x*0 = 60 dB; Δ*x* = 5 dB. D. Schematic showing the offset amplitude *y*0 parameter of the sigmoid fit (left panel) and cumulative distribution function of *y*0 for monotonic sound-increasing neurons at different laser powers of SST (middle panel) and VIP (right panel) neuronal activation. E. Schematic showing the amplitude range *y*range parameter of the sigmoid fit (left panel) and cumulative distribution function of *y*range for monotonic sound-increasing neurons at different laser powers of SST (middle panel) and VIP (right panel) neuronal activation. F. Schematic showing the midpoint *x*0 parameter of the sigmoid fit (left panel) and cumulative distribution function of *x*0 for monotonic sound-increasing neurons at different laser powers of SST (middle panel) and VIP (right panel) neuronal activation. G. Schematic showing the width Δ*x* parameter of the sigmoid fit (left panel) and cumulative distribution function of Δ*x* for monotonic sound-increasing neurons at different laser powers of SST (middle panel) and VIP (right panel) neuronal activation. H. Fit types (sigmoid in red, Gaussian in yellow, no fit in white) at one or both of the other laser powers for the non-SST neurons fit by a sigmoid function at no, medium and high laser powers of SST neuronal activation. The overlap between yellow and red (orange) indicates neurons fit by a sigmoid function at one laser power and a Gaussian function at the other laser power. I. Fit types (sigmoid in red, Gaussian in yellow, no fit in white) at one or both of the other laser powers for the non-VIP neurons fit by a sigmoid function at no, medium and high laser powers of VIP neuronal activation. The overlap between yellow and red (orange) indicates neurons fit by a sigmoid function at one laser power and a Gaussian function at the other laser power. J. Schematic of the mean significant changes to the response-level curve of a monotonic sound-increasing cell (gray line) upon SST (blue line) and VIP (green line) neuronal activation. For all panels, black, pink and red colors correspond to no laser power (0 mW/mm2), medium laser power (∼0.3 mW/mm2) and high laser power (∼3.5 mW/mm2), respectively (see Methods for power calibration). For all panels, the statistical test GLME was performed on the distributions at the three different levels of interneuron activation, with n.s. corresponding to non-significant, * corresponds to p<0.05, ** corresponds to p<0.01, *** corresponds to p<0.001. There are n=109, n=103 and n=64 sound-increasing monotonic cells fit at no, medium and high laser powers of SST neuronal activation, respectively. There are n=267, n=239 and n=269 sound-increasing monotonic cells fit at no, medium and high laser powers of VIP neuronal activation, respectively.

**FIGURE 5:**
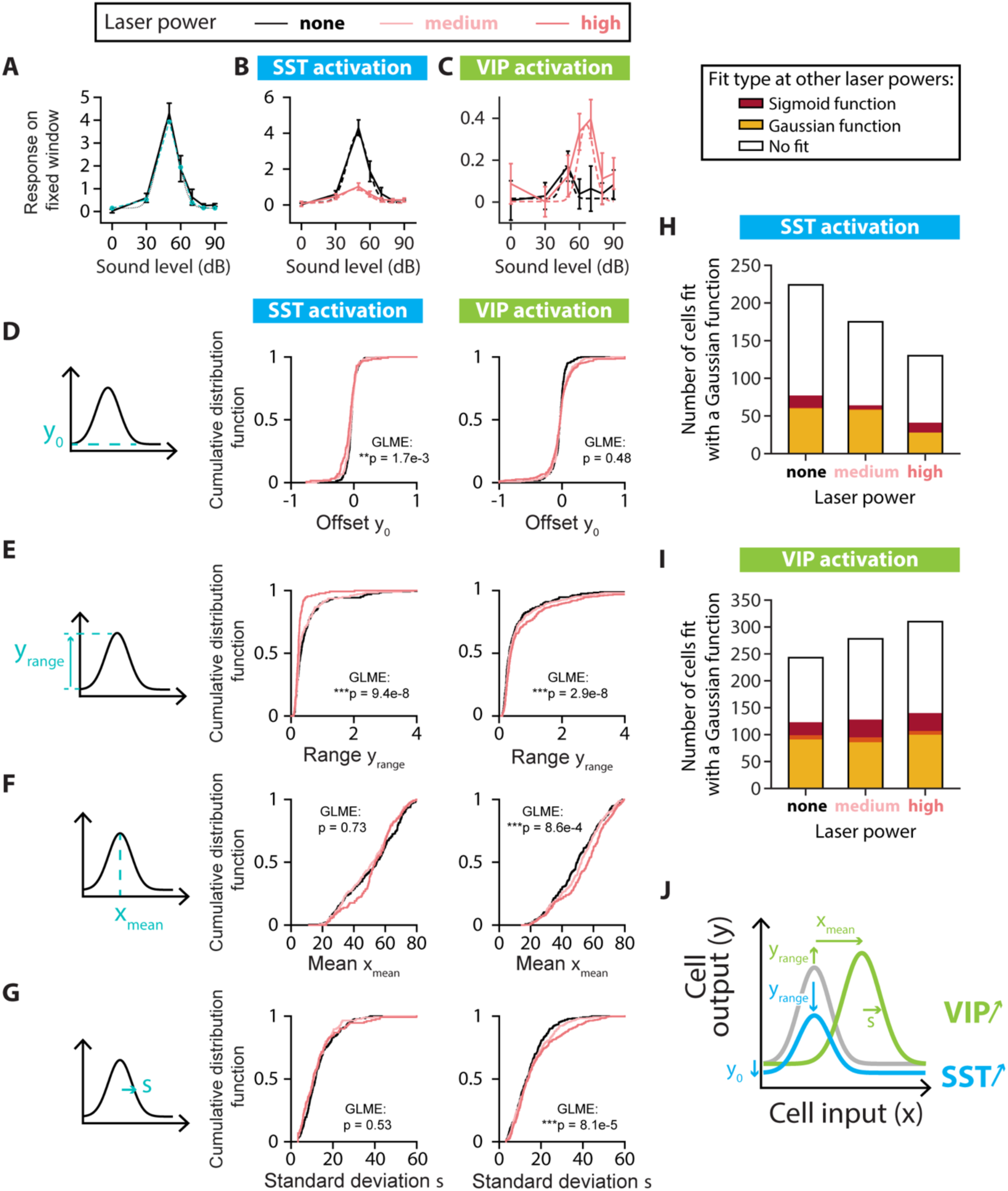
SINGLE-CELL FITS OF RESPONSE-LEVEL CURVES FOR NONMONOTONIC CELLS. A. Example neuron with a nonmonotonic response-level curve (solid black line) with a Gaussian fit estimated at the probed sound pressure levels (blue dashed line with circles) and the Gaussian function with the same parameters (dotted black line). The parameters for the Gaussian fit are: Offset amplitude *y*0 = 0.15; Range *y*range = 3.8; Mean *x*mean = 49 dB; Standard Deviation α = 9 dB. B. Example of changes to the response-level curve of a sound-increasing nonmonotonic neuron upon SST neuronal activation. Response-level curves (solid lines) are fit by Gaussian functions (dotted lines) at no (black lines) and high (red lines) laser powers of SST neuronal activation. The parameters for the Gaussian fit are, for no SST neuronal activation: *y*0 = 0.2; *y*range = 3.8; *x*mean = 49 dB; α= 9 dB; and for SST neuronal activation: *y*0 = 0.1; *y*range = 0.8; *x*mean = 47 dB; α= 11 dB. C. Example of changes to the response-level curve of a sound-increasing nonmonotonic neuron upon VIP neuronal activation. Response-level curves (solid lines) are fit by Gaussian functions (dotted lines) at no (black lines) and high (red lines) laser powers of VIP neuronal activation. The parameters for the Gaussian fit are, for no VIP neuronal activation: *y*0 = 0.0; *y*range = 0.2; *x*mean = 49 dB; α= 4 dB; and for VIP neuronal activation: *y*0 = 0.0; *y*range = 0.4; *x*mean = 66 dB; α= 7 dB. D. Schematic showing the offset amplitude *y*0 parameter of the Gaussian fit (left panel) and cumulative distribution function of *y*0 for nonmonotonic sound-increasing neurons at different laser powers of SST (middle panel) and VIP (right panel) neuronal activation. E. Schematic showing the amplitude range *y*range parameter of the Gaussian fit (left panel) and cumulative distribution function of *y*range for nonmonotonic sound-increasing neurons at different laser powers of SST (middle panel) and VIP (right panel) neuronal activation. F. Schematic showing the mean *x*mean parameter of the Gaussian fit (left panel) and cumulative distribution function of *x*mean for nonmonotonic sound-increasing neurons at different laser powers of SST (middle panel) and VIP (right panel) neuronal activation. G. Schematic showing the standard deviation α parameter of the Gaussian fit (left panel) and cumulative distribution function of α for nonmonotonic sound-increasing neurons at different laser powers of SST (middle panel) and VIP (right panel) neuronal activation. H. Fit types (sigmoid in red, Gaussian in yellow, no fit in white) at one or both of the other laser powers for the non-SST neurons fit by a Gaussian function at no, medium and high laser powers of SST neuronal activation. The overlap between yellow and red (orange) indicates neurons fit by a sigmoid function at one laser power and a Gaussian function at the other laser power. I. Fit types (sigmoid in red, Gaussian in yellow, no fit in white) at one or both of the other laser powers for the non-VIP neurons fit by a Gaussian function at no, medium and high laser powers of VIP neuronal activation. The overlap between yellow and red (orange) indicates neurons fit by a sigmoid function at one laser power and a Gaussian function at the other laser power. J. Schematic of the mean significant changes to the response-level curve of a nonmonotonic sound-increasing cell (gray line) upon SST (blue line) and VIP (green line) neuronal activation. For all panels, black, pink and red colors correspond to no laser power (0 mW/mm2), medium laser power (∼0.3 mW/mm2) and high laser power (∼3.5 mW/mm2), respectively (see Methods for power calibration). For all panels, the statistical test GLME was performed on the distributions at the three different levels of interneuron activation, with n.s. corresponding to non-significant, * corresponds to p<0.05, ** corresponds to p<0.01, *** corresponds to p<0.001. There are n=224, n=175 and n=130 sound-increasing nonmonotonic cells fit at no, medium and high laser powers of SST neuronal activation, respectively. There are n=243, n=278 and n=310 sound-increasing nonmonotonic cells fit at no, medium and high laser powers of VIP neuronal activation, respectively.

We first characterized the responses of sound-modulated cells that exhibited a monotonic response-level curve by fitting this curve for individual cells at different levels of laser power (Figure 4A-C). Out of the four sigmoidal fit parameters (Figure 4D-G, middle panels), only the midpoint of the sigmoid fit exhibited a significant change upon SST neuronal activation from 58 dB at no laser power to 73 dB at high laser power (median values, n=109; 103 and 64 neurons for no, medium and high laser power, respectively; offset: *p*_laser_=0.41; range: *p*_laser_=0.22; midpoint: \*\**p*_laser_=8.6e-3; width: *p*_laser_=0.99; GLME, Figure 4D-G, middle panels). With VIP activation, only the range of the sigmoid fit showed a significant increase (n=267, 239 and 269 neurons for no, medium, and high laser power, respectively; offset: *p*_laser_=0.081; range: \*\**p*_laser_=1.9e-3; midpoint: *p*_laser_=0.38; width: *p*_laser_=0.57; GLME, Figure 4D-G, right panels). Among the non-SST neurons fit by a sigmoidal function, two thirds of the cells were fit at a single laser power of SST neuronal activation (n=75, 60 and 40 neurons for no, medium and high laser power, respectively, Figure 4H) and around a tenth of the neurons switched monotonicity with SST neuronal activation (Gaussian fit at other laser powers for n=10, 21 and 8 neurons at no, medium and high laser power, respectively). Among the non-VIP neurons fit by a sigmoidal function, around 40% of the cells were fit at a single laser power of VIP neuronal activation (n=108, 88 and 109 neurons for no, medium and high laser power, respectively, Figure 4I) and around 15% of the neurons switched monotonicity with VIP neuronal activation (Gaussian fit at other laser powers for n=49, 35 and 39 neurons at no, medium and high laser power, respectively). Thus, SST neuronal activation leads to monotonic response-level functions that are shifted rightwards towards higher sound pressure levels, leading to responses to a narrower range of sounds at higher sound pressure levels. VIP activation expanded the neuronal response-level curves upwards, leading to responses to a broader range of sound pressure levels (Figure 4J).

We then characterized the responses of sound-modulated cells that exhibited a nonmonotonic response-level curve by fitting this curve for individual cells at different levels of laser power (Figure 5A-C). Whereas the mean and standard deviation of the Gaussian fit remained unchanged (mean: *p*_laser_=0.73; standard deviation: *p*_laser_=0.53; GLME, Figure 5F-G, middle panels), the offset and the range of the Gaussian fit decreased with SST neuronal activation (n=224, 175 and 130 neurons for no, medium and high laser power, respectively; offset: \*\**p*_laser_=1.7e-3; range: \*\*\**p*_laser_=9.4e-8; GLME, Figure 5D-E, middle panels). The offset of response did not change with VIP neuronal activation (offset: *p*_laser_=0.48, GLME, Figure 5D, right panel), whereas the range increased significantly as well as the Gaussian mean and standard deviation (n=243, 278 and 310 neurons for no, medium and high laser power, respectively; range: \*\*\**p*_laser_=2.9e-8; mean: \*\*\**p*_laser_=8.6e-4; standard deviation: \*\*\**p*_laser_=8.1e-5; GLME, Figure 5E-G, right panels). Among the non-SST neurons fit by a Gaussian function, two thirds of the cells were fit at a single laser power of SST neuronal activation (n=147, 111 and 89 neurons for no, medium and high laser power, respectively, Figure 5H) and less than a tenth of the neurons switched monotonicity with SST neuronal activation (sigmoidal fit at other laser powers for n=17, 6 and 13 neurons at no, medium and high laser power, respectively). Among the non-VIP neurons fit by a Gaussian function, half of the cells were fit at a single laser power of VIP neuronal activation (n=120, 150 and 170 neurons for no, medium and high laser power, respectively, Figure 5I) and around 15% of the neurons switched monotonicity with VIP neuronal activation (sigmoidal fit at other laser powers for n=32, 42 and 40 neurons at no, medium and high laser power, respectively). Thus, SST neuronal activation led to nonmonotonic response-level functions that were shifted downwards with a decreased range of responses, leading to responses above noise level to a narrower range of sound pressure levels. VIP neuronal activation shifted the neuronal response-level curves rightwards with an increased range of response, leading to increased peak responses at higher sound pressure levels (Figure 5J).

These results demonstrated that SST neuronal activation promoted a more localist representation of sound pressure level at the population scale by eliciting responses above noise level over a narrower range of sound pressure levels, and increasing separation between the sound pressure levels covered by monotonic and nonmonotonic neurons. By contrast, VIP neuronal activation promoted a more distributed representation of sound pressure level by broadening response-level curves of single neurons and increasing the overlap between the sound pressure levels covered by monotonic and nonmonotonic neurons.

Could the changes to the fitted parameters of sound-modulated cells (Figure 4 and Figure 5) explain the differential effect of SST and VIP neuronal activation on the separation angle and length between mean population vectors to different sound pressure levels (Figure 3)? To answer this question, we constructed a qualitative model including a monotonic and a nonmonotonic cell, with response-level fit parameters for no and high laser power taken as the mean parameters from the data in Figure 4 and Figure 5 for no and high laser power (Figure 6A-B). With this simple two-cell population, we could qualitatively reproduce the increase in separation angle upon SST neuronal activation and the decrease in separation angle upon VIP activation over the range of sound pressure levels we tested (Figure 6D-F), and similarly the decrease in vector length upon SST neuronal activation and the increase in vector length upon VIP (Figure 6G-I). We could also model different regimes of multiplicative (or divisive) and additive (or subtractive) effects of SST or VIP neuronal activation on monotonic and nonmonotonic neurons (Figure 6J-K). Thus, the changes to the fit parameters observed in Figures 4-5 can explain the change in representation of sound pressure levels observed in Figure 3.

**FIGURE 6:**
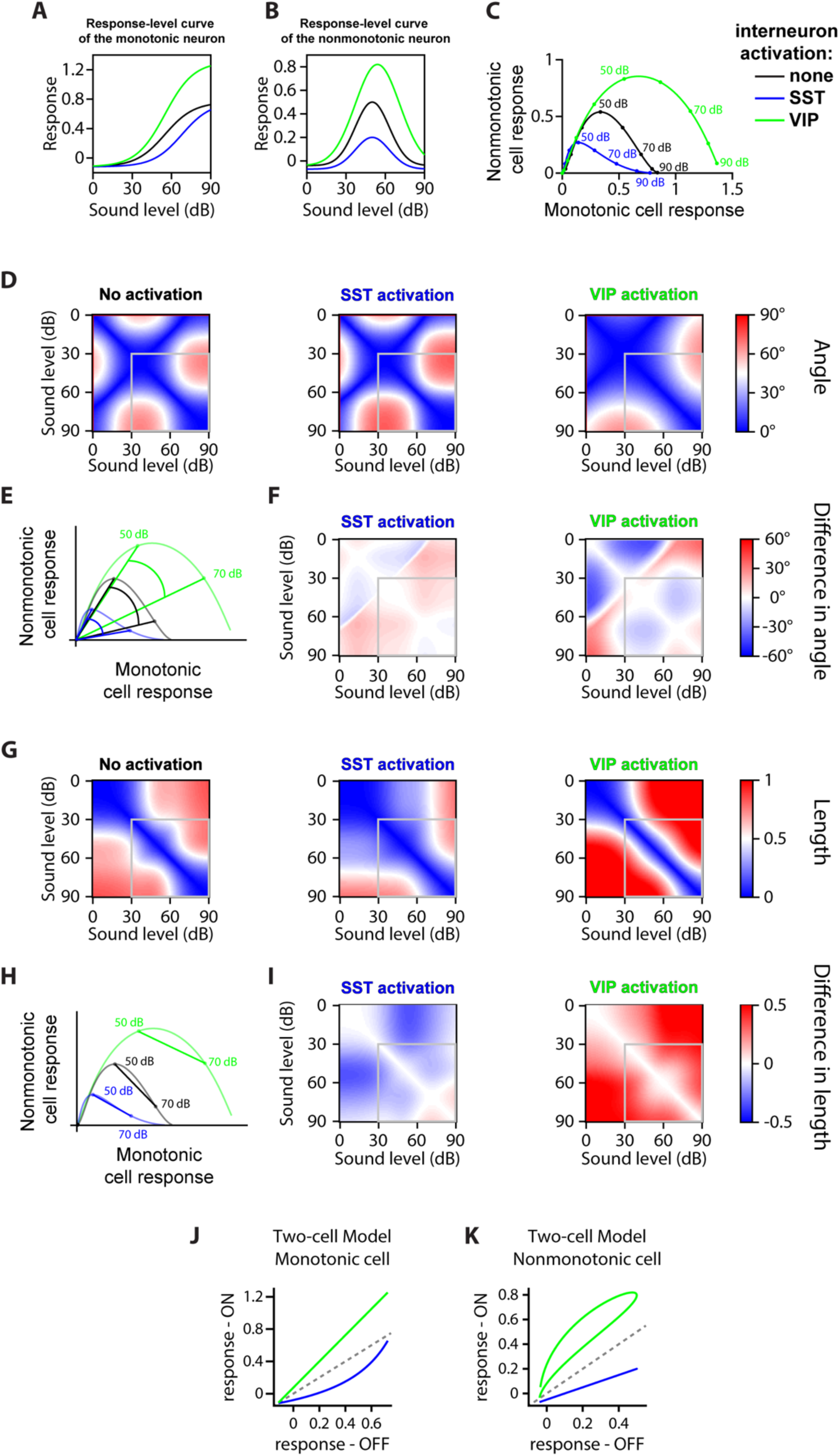
TWO-CELL MODEL. A. Response-level curve of the monotonic cell with parameters taken as the mean parameters from Figure 4 at no and high laser power. The parameters for the sigmoid curve with no interneuron activation (black) are: Offset amplitude y_0_ = -0.12; Range y_range_ = 0.88; Midpoint x_0_ = 55 dB; Width Dx = 11 dB. Upon SST neuronal activation (blue), all parameters remain constant except for: Midpoint x_0_^SST^ = 68 dB; Upon VIP neuronal activation (green), all parameters remain constant except for: Range y_range_^VIP^ = 1.43. B. Response-level curve of the nonmonotonic cell with parameters taken as the mean parameters from Figure 5 at no and high laser power. The parameters for the Gaussian curve with no interneuron activation (black) are: Offset amplitude y_0_ = -0.04; Range y_range_ = 0.54; Mean x_mean_ = 50 dB; Standard Deviation s = 13 dB. Upon SST neuronal activation (blue), all parameters remain constant except for: Offset amplitude y_0_^SST^ = -0.07; Range y_range_^SST^ = 0.27; Upon VIP neuronal activation (green), the offset amplitude remains constant, and: Range y_range_^VIP^ = 0.86; Mean x_mean_^VIP^ = 54 dB; Standard Deviation s^VIP^ = 17 dB. C. Trajectory of the population’s response from 0dB to 90dB in the neural space, with the response of the monotonic cell on the x-axis and the response of the nonmonotonic cell on the y-axis. The response of both cells at 0dB has been subtracted from the curves, thus the dots at the (0,0) coordinate are the response to 0dB, and the end of the curves on the right indicate the response to 90dB. The trajectories are computed from 0 dB to 90dB with 1dB increments, and circles on a line represent 10dB increments from 0dB to 90dB. D. Confusion matrix of the separation angle between population responses to each sound and laser power from silence at a given laser power, for no (left), SST (middle) and VIP (right) neuronal activation. Sound pressure level is in 1dB increments, and the gray box indicates the sound levels sampled in the experiments (Figure 3D, and G) E. Schematic in the neural space (see panel C) of the angle between 50 dB and 70dB when there is no (black), SST (blue) or VIP (green) neuronal activation, starting from the population’s response to silence for each case of (or lack of) interneuron activation. The angle is greatest when SST neurons are activated, and smallest when VIP neurons are activated. F. Confusion matrix of the difference in separation angle from SST (left) or VIP (right) neuronal activation to no interneuron activation, with the angles calculated as in (D). Sound level is in 1dB increments, and the gray box indicates the sound pressure levels sampled in the experiments (Figure 3F and I). The mean angle difference for SST neuronal activation is, over 1-90dB: + 3.6° and over 30-90dB: + 3.7°; for VIP activation, over 1-90dB: − 4.0°, and over 30-90dB: − 4.1°. G. Confusion matrix of the length of the population vector between each sound pressure level at a given laser power, for no (left), SST (middle) and VIP (right) neuronal activation. Sound pressure level is in 1dB increments, and the gray box indicates the sound pressure levels sampled in the experiments (Figure 3J and M). H. Schematic in the neural space (see panel C) of the vector length between 50 dB and 70dB when there is no (black), SST (blue) or VIP (green) neuronal activation. The vector length is greatest when VIP neurons are activated, and smallest when SST neurons are activated. I. Confusion matrix of the difference in vector length from SST (left) or VIP (right) neuronal activation to no interneuron activation, with the lengths calculated as in (G). Sound pressure level is in 1dB increments, and the gray box indicates the sound pressure levels sampled in the experiments (Figure 3I and L). The mean length difference for SST neuronal activation is, over 1-90dB: − 0.12 a.u. and over 30-90dB: − 0.07 a.u.; for VIP neuronal activation, over 1-90dB, + 0.27 a.u. and over 30-90dB: + 0.21 a.u. J. Response of the monotonic cell with laser activation versus no laser activation. SST neuronal activation shows a divisive regime, a subtractive regime, a combination of divisive and subtractive or multiplicative and subtractive regimes depending on the range of responses sampled. VIP neuronal activation shows a multiplicative regime or an additive and multiplicative regime depending on the range of responses sampled. K. Response of the nonmonotonic cell with laser activation versus no laser activation. SST neuronal activation shows a combination of divisive and subtractive regime. VIP neuronal activation shows a multiplicative regime, and additive regime or an additive and multiplicative regime depending on the range of responses sampled. For all panels, black indicates no interneuron activation, blue indicates SST neuronal activation and green indicates VIP neuronal activation.

Overall, when neither SST or VIP neurons are activated, the neuronal population encoded different sound pressure levels using two strategies: the identity of the responsive cells (different cells respond to different sound pressure levels, discrete encoding of sound pressure level) and the strength of the neuronal response (continuous encoding of sound pressure level). When SST neurons are activated, the neuronal population shifts towards a more localist representation of sound pressure level. Specifically, the encoding of sound pressure level relies more on the identity of the responsive cells and less on the magnitude of response: there is less overlap between populations of cells responding to different sound pressure levels, but the strength of response is similar for all sound pressure levels. This can be explained with the narrower bandwidths of response, albeit of reduced magnitude, of both monotonic and nonmonotonic sound-increasing cells with SST neuronal activation. By contrast, when VIP neurons are activated, the neuronal population shifts towards a more distributed representation of sound pressure level. Specifically, the encoding of sound pressure level by the magnitude of the neuronal response is enhanced, while the representation by different cell groups declines: the neuronal responses are of higher magnitude and over a higher range, but there is more overlap between neurons responding to different sound pressure levels. This can be explained with the larger and broader responses of monotonic and nonmonotonic sound-increasing neuronal responses with VIP neuronal activation.

## DISCUSSION

Within the brain, neurons form intricate networks, which represent sensory information. A sensory stimulus, such as a specific sound or a visual image, elicits activity in a subset of neurons in a network. A neuronal network can use a multitude of codes to represent information. A stimulus can be encoded discretely with a localist, pattern-separated representation, in which a specific group of neurons represents a specific stimulus, and different stimuli elicit activity in different groups of neurons (Figure 7A). Such localist representations have the advantage of discreteness: they can separate stimuli in different categories. Alternatively, in a distributed representation, stimulus-evoked activity can be distributed across the network, such that the relative activity of neurons within a group represent different stimuli (Figure 7A-B). An example of a distributed representation is a rate code, in which the firing rate of the active neurons represent a continuously varying stimulus feature, such as intensity or sound location (Belliveau et al., 2014). Distributed representations have the advantage of invariance: a small change in stimulus parameter will elicit a small variation in the neuronal response. Neuronal population responses have been measured at various positions along the localist to distributed representation spectrum across many features and areas (such as memory: (Wixted et al., 2014), sound (Hromádka et al., 2008), sound localization (Lesica et al., 2010; Belliveau et al., 2014), vision (Christensen and Pillow, 2022)) and can change dynamically along the spectrum (Kato et al., 2015; Honey et al., 2017; Kuchibhotla et al., 2017). A defining feature of the auditory cortex is sparse coding (DeWeese et al., 2003), which can lead to both distributed and localist representations. Along the auditory pathway, the coexistence of neurons with monotonic and nonmonotonic response-level curves indicates that sound pressure level is represented by both localist and distributed codes (Schreiner et al., 1992; Wu et al., 2006; Tan et al., 2007; Watkins and Barbour, 2011). More generally, stimulus representation within neuronal networks is mixed between local and distributed codes, in so-called sparse distributed representations, with both the level of activity and identity of activated neurons encoding the stimulus (Figure 7A) (Rolls and Tovee, 1995; Vinje and Gallant, 2000; Hromádka et al., 2008; Wixted et al., 2014). Based on the environmental and behavioral demands, it may be beneficial for neuronal representations to shift dynamically towards a more localist or a more distributed representation of a stimulus feature.

**FIGURE 7:**
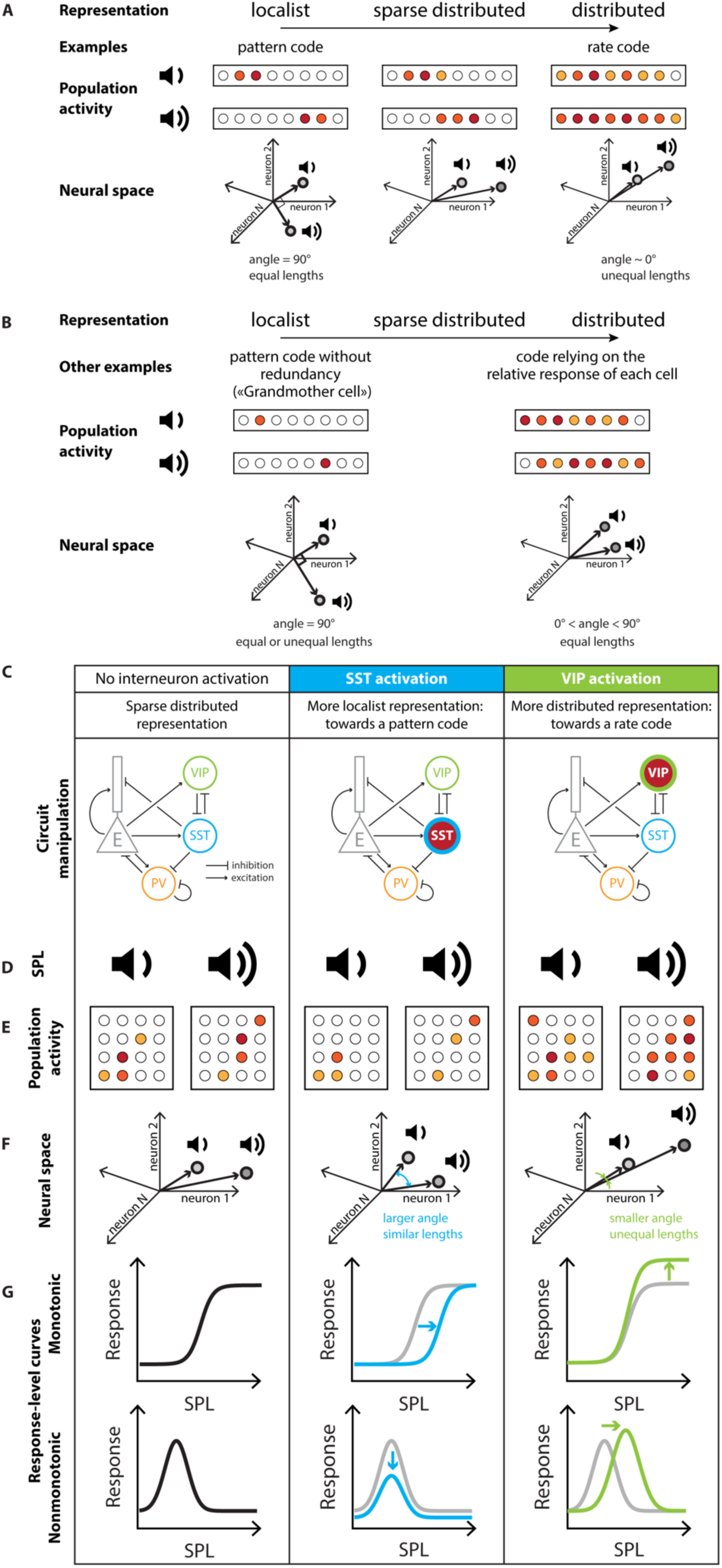
PREDICTIONS FOR TESTING FOR DIFFERENTIAL CONTROL OF LOCALIST AND DISTRIBUTED REPRESENTATIONS OF SOUNDS BY INHIBITORY NEURONS. A. A schematic of example localist versus distributed neuronal codes. B. Localist versus distributed representations can be implemented in many ways. An example of localist code is the pattern code without redundancy (known as the “Grandmother cell”, (Bowers, 2009)). An example of distributed code is a code relying only on the relative response of each cell and not the magnitude of response of the population vector. C. Neuronal circuit manipulation: Optogenetic stimulation of SST and VIP neurons in Auditory Cortex (simplified connectivity circuit). D. Noise bursts at different sound pressure levels (SPL) were presented to an awake mouse. E. For each sound pressure level, some neurons responded (filled circles) while many cells didn’t respond (empty circles). F. The response to different sound pressure levels can be described in the neuronal space by the separation angle between the mean population vectors and the length separating them. G. Changes to the representation of sound pressure level at the population level are implemented through changes to the response-level curve of each neuron. Gray: baseline; blue: response transformation with SST neuronal activation; green: response transformation with VIP neuronal activation.

Our results suggest that distinct inhibitory neurons in the auditory cortex affect population neuronal response code by differentially shifting the responses toward a distributed or a localist representations. Previous work found that SST neuronal activation decreases the activity of the neuronal population to sounds (Phillips and Hasenstaub, 2016; Natan et al., 2017), and leads to a rightward shift of monotonic response-level curves (Wilson et al., 2012), while VIP neuronal activation, through a disinhibitory circuit, increases the activity of the population (Pfeffer et al., 2013; Pi et al., 2013; Zhang et al., 2014). We found that activation of SST neurons, led to a sparser, more localist representation (Figure 2D-E), where sound pressure level is encoded in discrete steps by distinct groups of neurons (Figure 3E) with a similar low strength (Figure 3K). By contrast, activation of VIP neurons, led to a more distributed representation (Figure 2K-L), with more overlap between the cell populations responding to different sound pressure levels (Figure 3H): sound pressure level is encoded continuously by varying the strength of response of a large group of neurons (Figure 3N). These shifts in representation are implemented at the single-neuron level through changes to the response-level function of monotonic (Figure 4) and nonmonotonic neurons (Figure 5). SST neuronal activation shifts the response-level curves of sound-modulated neurons by further separating the sound pressure levels that monotonic and nonmonotonic neurons represent (Figure 4I and 5I). With VIP neuronal activation, the changes to the response-level curves of sound-modulated neurons allow for stronger responses and more overlap in bandwidth between monotonic and nonmonotonic neurons (Figure 4J and 5J).

As distinct inhibitory populations can be recruited during different behaviors, their ability to transform the neuronal code can be advantageous to distinct behaviors and computations. Localist versus distributed representations, modulated by the relative strength of global (SST) versus local (PV or VIP) inhibition respectively, may provide support for neuronal computations such as discreteness versus invariance (Kuchibhotla and Bathellier, 2018), segmentation versus concatenation (Haga and Fukai, 2021), integration of bottom-up versus top-down information (Honey et al., 2017; Hertäg and Sprekeler, 2019). The two types of interneurons receive neuromodulatory inputs and may dynamically change the network’s state towards one or the other type of representation for a given task: SST neuronal activation may help with tasks requiring focused attention such as discriminating different stimuli (Lee and Middlebrooks, 2011) or detecting in noise (Lakunina et al., 2022), by sharpening tuning, decreasing the activity for non-relevant stimuli, and enhancing the information-per-spike (Phillips and Hasenstaub, 2016). In contrast, VIP neuronal activation may help with tasks requiring receptive-attention such as active sensing (Gentet et al., 2012) or detecting small stimuli by amplifying weak signals (Millman et al., 2020), increasing detectability (Cone et al., 2019) without increasing the stimulus-response mutual information (Bigelow et al., 2019).

Our results expand on the results of previous studies that measured more general effects of interneuron modulation. The changes to the response-level curves (Figures 4 and 5) we measured upon SST or VIP neuronal activation can explain how the frequency-response functions of neurons in AC change with interneuron modulation. The excitatory and inhibitory inputs to pyramidal cells of AC are frequency-tuned (Isaacson and Scanziani, 2011; Li et al., 2013; Kato et al., 2017), and intracortical inhibition further shapes the tuning (Wu et al., 2008). From the response-level curves, we can plot the response with interneuron activation versus without and thus predict within which range of amplitudes we can expect multiplicative or additive effects to the frequency tuning curve. Previous studies have shown that SST neuronal activation leads to either subtractive, divisive or a combination of both subtractive and divisive effects on the frequency tuning curve (Wilson et al., 2012; Seybold et al., 2015; Phillips and Hasenstaub, 2016). Our data are consistent with these results: for monotonic neurons, SST neuronal activation can lead to divisive and/or subtractive effects at a low range of sound amplitudes and for nonmonotonic neurons, SST neuronal activation leads to both divisive and subtractive effects (Figure 6J-K). Previous studies have also shown that VIP leads to an additive shift in the frequency tuning curve (Pi et al., 2013; Bigelow et al., 2019) and similarly SST inactivation leads to a multiplicative or additive shift mostly (Phillips and Hasenstaub, 2016). Our data can explain these results as well, with multiplicative effects for monotonic neurons and a range of additive and multiplicative effects for nonmonotonic neurons (Figure 6J-K). One component that can contribute to a change in representation is a change in the noisiness of the responses. Indeed, changes in the signal-to-noise ratio (SNR) could drive populations to appear as more localist or distributed in their coding. The change in SNR may be one of the components explaining how the change in representation is implemented, however it is not the sole factor as assessed by the decoding accuracy (Figure 2F and M).

Nonmonotonic neurons in AC either can have their nonmonotonicity inherited from the nonmonotonic excitatory input into those cells while the monotonic inhibitory input, which shows a peak in delay at the cell’s best pressure level, sharpens the nonmonotonicity (Wu et al., 2006) or can be constructed *de novo* with an imbalance between excitatory and inhibitory inputs (Wu et al., 2006; Tan et al., 2007). This means that for some of the nonmonotonic cells we recorded from, the input into the cell does not covary with the sound pressure level, and so for these cells, we are not assessing the input-output function of the cell through the response-level curve, rather how inhibition further shapes the already intensity-tuned input. From our experiments, we observe that SST neuronal activation decreases the range of responses of nonmonotonic neurons but does not change the sound pressure level of peak response nor the width of responses (Figure 5). This may indicate that SST neuronal activation does not change the timing of the inhibitory input into the nonmonotonic cells, but rather the overall strength of inhibition across all sound pressure levels. In contrast, VIP neuronal activation leads to a shift of the sound pressure level of peak response towards higher levels, along with a broadening of the response and an increase in the range of response (Figure 5). This could simply be explained if VIP neuronal activation changes the timing of the inhibition, with the delay between excitatory and inhibitory inputs peaking at a higher sound pressure level.

In our sample, the relative proportion of nonmonotonic versus monotonic neurons is higher than would be expected from the literature. We note that the number of neuronal responses that we were able to fit are likely an underestimate of the truly monotonic or non-monotonic neurons in the population, as we used stringent selection criteria. The relatively high proportion of non-monotonic neurons may be due to inclusion of the non-primary region VAF, which has previously been shown to have a higher proportion of nonmonotonic neurons than A1 (Wu et al., 2006; Polley et al., 2007). Furthermore, we used a mouse line in which the hearing loss mutation in Cdh23 commonly found in C57B6 mice is corrected, thus our mice may have lower detection thresholds than the mice in previous studies, leading to a larger observed proportion of nonmonotonic neurons tuned to lower intensities. Additionally, because of the time course of GCaMP, we are not distinguishing between onset and offset responses, and they may be integrated in this window. These estimates contribute to the ongoing discussion of the differences in monotonicity of responses between the non-human primates (Gao and Wang, 2019) and rodents. It is plausible that our current setup, with imaging performed in awake mice rather than under anesthesia, allows us to sample the responses in a more accurate fashion than previous studies.

In our analysis, the majority of the neurons remained monotonic or non-monotonic between laser conditions, with only 10-15% switching monotonicity (Figures 4H-I and 5H-I). Additionally, the ratio between the neurons which we identify as monotonic or non-monotonic was generally preserved across the laser conditions. A significant fraction of neurons were fit at a single laser power of SST or VIP neuronal activation. SST and VIP neuronal activation thus elicited responses in new pools of neurons for each laser, however with more consistency between laser powers upon VIP neuronal activation than upon SST neuronal activation. Because of the stringent fitting criteria, we believe that our fitting procedure underestimates the true fraction of monotonic or non-monotonic neurons. Nonetheless, we are able to compare the fits across conditions because the criteria and fraction of well-fitted responses remain the same. Therefore, the change in representation (localist or distributed) likely relies on changes to the response-level curves rather than on changes in the proportion of monotonic and nonmonotonic neurons.

A limitation of our study is that we measured the response only from a subset of neurons from layer 2/3 while broadly stimulating many SST or VIP neurons across different cortical layers. However, neurons across the cortical column may perform additional computations in other layers, which would be important to record in future experiments. Our sample ended up including a relatively low number of SST-positive and VIP-positive neurons. Their sound intensity responses largely trended the mean recorded responses and did not differ strongly from each other. From the literature, we were expecting VIP neurons to be selective for low dB sounds (Mesik et al., 2015). Similarly, in the visual cortex, VIP neurons prefer low contrast whereas SST neurons prefer high contrast visual stimuli (Millman et al., 2020). Here, SST and VIP neurons have similar tuning properties with the non-SST and non-VIP neurons within their sessions, but with optogenetic stimulation, sound modulation becomes weaker. Future study should focus on the responses within the SST and VIP neuronal populations across layers. Furthermore, whereas we combined imaging sessions across the auditory cortex, future studies should also examine differences in function of inhibitory neurons across the different primary and non-primary auditory areas.

A related caveat in interpreting our results is that optogenetic activation of inhibitory neurons may differ across samples, and can potentially drive higher activity level of excitatory or inhibitory neurons than physiological levels, saturating the responses. To mitigate this limitation, we included multiple activation levels in each of the imaging sessions by modulating the strength of the laser between high and low levels. This allowed within sample comparison of activity patterns, rather than comparisons of absolute changes across multiple imaging sessions. Furthermore, by comparing the activity levels of imaged neurons between low and high laser intensities, we ensured that the modulation of activity was not saturating due to the laser.

An additional caveat is that the opsin-expressing cells may be depolarized at baseline due to off-target laser stimulation during two-photon imaging (Forli et al., 2018). In SST-Cre mice, the depolarization of SST neurons would lead monotonic non-SST neurons to shift their range of responsiveness towards higher sound pressure levels (Figure 4), while nonmonotonic non-SST neurons still respond to the same best sound level. This would lead to a proportionally larger response to the lower sound pressure levels (covered mainly by nonmonotonic neurons) than to the high sound pressure levels, as we observe in Figure 2C. In VIP-Cre mice, the shape of the population response-level curve (Figure 2J) is similar to that in SST-Cre mice. The depolarization of VIP neurons at baseline may lead to an increase in the amplitude range of nonmonotonic neurons combined with a slight increase in the number of nonmonotonic neurons, which may result in similar changes to the population response-sound level curve compared to the curve in the control.

Another potential caveat in interpreting the data is the while we were able to identify a subset of neurons as SST or VIP positive, the analysis combined multiple types of neurons as non-SST or non-VIP neurons, and for some neurons the tdTomato expression might not have been strong enough for detection as SST or VIP positive. The population of unlabelled neurons may include SST neurons explaining the increase in population response for non-SST neurons in response to high laser power and no sound (Figure 2C). SST neurons also suppress PV neurons (Pfeffer et al., 2013), and at high laser power, in combination with firing rate saturation and indirect disinhibition of excitatory neurons via PV neurons, the overall activity might be increasing in the absence of sound. A targeted experimental approach which would allow either in vivo or post-hoc identification of imaged neuronal types would allow to further distinguish response parameters among different types of inhibitory and excitatory neurons (Kerlin et al., 2010; Khan et al., 2018). Furthermore, future studies inactivating these inhibitory neurons would further complement and extend our findings (Phillips and Hasenstaub, 2016).

In our experiments, we tested the results of SST or VIP neuronal activation on the neuronal representation of sound pressure level, and an important next step will be to investigate whether the changes in neuronal representation of sound pressure level correlate with behavioral effects. One approach would be to image the activity of SST neurons while a mouse is engaged in a task that may require SST neuronal activation, or similarly VIP neurons. SST neurons may be involved in tasks leading to a sharpening in the neuronal tuning properties, or to a filtering out of irrelevant stimuli through the overall decrease in firing rate, such as discrimination tasks and signal-in-noise tasks (Otazu et al., 2009; Lee and Middlebrooks, 2011; Kuchibhotla et al., 2017; Christensen et al., 2019; Lakunina et al., 2022). VIP neurons may be involved in tasks requiring amplification of weak signals, such as a detection at threshold task or active sensing (Fritz et al., 2003; Gentet et al., 2012; Bennett et al., 2013; Kato et al., 2015; Cone et al., 2019; Millman et al., 2020). A second approach would be to measure how SST or VIP neuronal activity changes the performance of a mouse engaged in a task. In a detection task, we may expect that detection thresholds increase with SST neuronal activation and decrease with VIP neuronal activation, as seen in the visual cortex (Cone et al., 2019). In a discrimination task or detection of sounds in background task, we may expect SST neuronal activation to increase the performance, while VIP neuronal activation may decrease or not affect performance: SST neuronal inactivation decreases performance in the detection of sounds in background noise (Lakunina et al., 2022), and at the neuronal level, VIP neuronal activation decreases “encoding efficiency” (Bigelow et al., 2019) while SST neuronal activation may increase it (Phillips and Hasenstaub, 2016).

It should be noted that the dichotomy between the functional roles of SST and VIP neurons might not be so clear cut: VIP and SST neurons may cooperate to simultaneously amplify relevant stimuli and filter out irrelevant stimuli, respectively, or VIP neurons may be more active for weak stimuli and SST neurons for loud stimuli (Zhang et al., 2014; Mesik et al., 2015; Karnani et al., 2016; Kuchibhotla et al., 2017; Dipoppa et al., 2018a; Millman et al., 2020). For example, activation of the cingulate cortex can elicit spiking activity in PV, SST and VIP neurons, with SST neurons contributing to surround suppression, and VIP neurons to facilitation of the center of the receptive field (Zhang et al., 2014); cholinergic modulation depolarizes multiple types of inhibitory neurons, including SST and VIP neurons (Kuchibhotla et al., 2017); and locomotion increases activity in VIP neurons primarily for small stimuli, but in SST neurons for large stimuli (Dipoppa et al., 2018b). VIP neurons inhibit SST neurons, creating attentional spotlights through targeted disinhibition (Karnani et al., 2016). Therefore, examination of the function of SST and VIP neurons in complex behaviors needs to take the circuit mechanisms into account. Interestingly, the connectivity between cortical neurons is largely conserved across layers, primary and non-primary cortices, sensory and non-sensory areas (Douglas and Martin, 2004; Markram et al., 2004; Yu et al., 2019; Campagnola et al., 2022), with SST and VIP neurons mutually inhibiting each other as a common motif: perhaps the change in representation we observe with varying sound level pressure upon SST or VIP neurons extends to other stimulus features, sensory and non-sensory alike.

## CONFLICT OF INTEREST

The authors declare no competing interests.

## ACKNOWLEDGEMENTS

The authors thank Dr. Anna Schapiro and members of the Geffen laboratory for helpful discussions and advice. This work was supported by NIDCD R01DC015527, R01DC014479, NINDS R01NS113241 to MNG and NIDCD K99DC019504 to KCW.

## Notes

https://doi.org/10.5061/dryad.t1g1jwt6d

### Competing Interest Statement

The authors have declared no competing interest.

### Summary of Updates

Minor additional analysis and reorganization of the manuscript to only have 7 figures (and no supplementary figures).

