## Extended Figures for "“Distinct inhibitory neurons differently shape neuronal codes for sound intensity in the auditory cortex”" (from another version of this preprint: version 4, 2024-07-06)

### Extended Figures: Legends and statistics

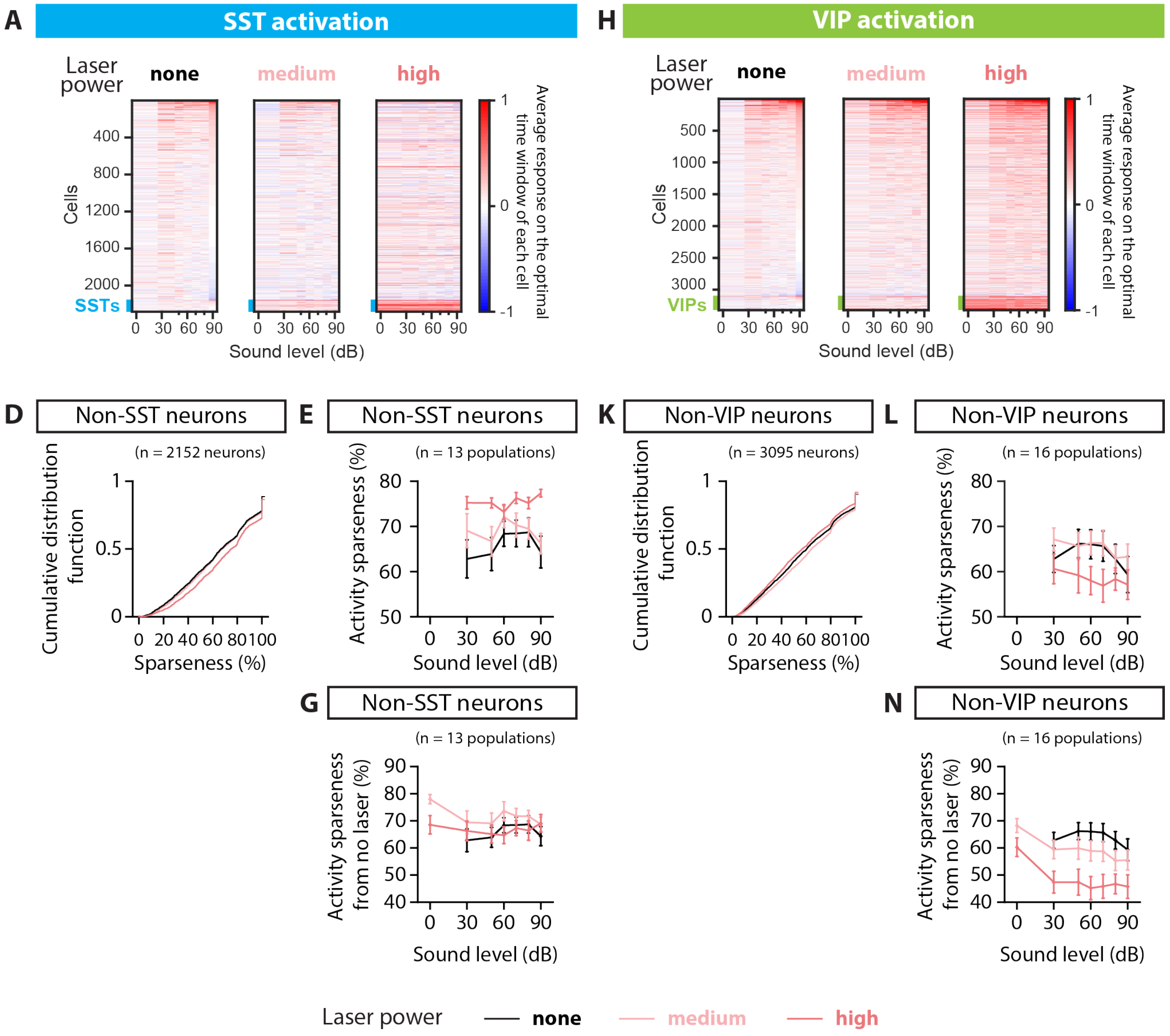


Extended Figure 2.1. Panels A, D, E, G, H, K, L, N of Figure 2 computed on a fixed [0 1] sec window instead of the optimal window.

| **EXTENDED FIGURE 2.1** | | | | | | |
|---|---|---|---|---|---|---|
| Sparseness with SST activation | Ext. Fig 2.2 D | None: 2020<br>Med: 2019<br>High: 1980 | GLME | $t_{laser}$=6.01<br>DF = 5676 | *****$p_{laser}$=1.9e-9** | $h_{laser}^2$=5.9e-3 |

| | | | | | | |
|---|---|---|---|---|---|---|
| Activity sparseness with SST activation | Ext. Fig 2.2 E | 13 populations | GLME | $t_{laser}$=3.06<br>$t_{sound}$=0.93<br>$t_{laser:sound}$=-0.56<br><br>DF = 230 | **$^{**}p_{laser}$=2.5e-3**<br>$p_{sound}$=0.35<br>$p_{laser:sound}$=0.57 | $h_{laser}^2$=0.19<br>$h_{sound}^2$=1.4e-2<br>$h_{laser:sound}^2$=1.2e-2 |
| Activity sparseness from 0dB and no laser power with SST activation | Ext. Fig 2.2 G | 13 populations | GLME | $t_{laser}$=0.31<br>$t_{sound}$=0.74<br>$t_{laser:sound}$=-0.23<br><br>DF = 230 | $p_{laser}$=0.76<br>$p_{sound}$=0.46<br>$p_{laser:sound}$=0.82 | $h_{laser}^2$=2.4e-3<br>$h_{sound}^2$=8.3e-3<br>$h_{laser:sound}^2$=2.1e-3 |
| Sparseness with VIP activation | Ext. Fig 2.2 K | None: 2997<br>Med: 3002<br>High: 3023 | GLME | $t_{laser}$=-3.11<br><br>DF = 8442 | **$^{**}p_{laser}$=1.8e-3** | $h_{laser}^2$=9.3e-4 |
| Activity sparseness with VIP activation | Ext. Fig 2.2 L | 16 populations | GLME | $t_{laser}$=-1.46<br>$t_{sound}$=-0.97<br>$t_{laser:sound}$=5.1e-4<br><br>DF = 284 | $p_{laser}$=0.14<br>$p_{sound}$=0.33<br>$p_{laser:sound}$=0.99 | $h_{laser}^2$=3.1e-2<br>$h_{sound}^2$=8.6e-3<br>$h_{laser:sound}^2$=6.0e-9 |
| Activity sparseness from 0dB and no laser power with VIP activation | Ext. Fig 2.2 N | 16 populations | GLME | $t_{laser}$=-5.54<br>$t_{sound}$=-1.45<br>$t_{laser:sound}$=0.58<br><br>DF = 284 | **$^{***}p_{laser}$=7.0e-8**<br>$p_{sound}$=0.15<br>$p_{laser:sound}$=0.56 | $h_{laser}^2$=0.22<br>$h_{sound}^2$=1.2e-2<br>$h_{laser:sound}^2$=4.7e-3 |

Extended Table 2.1: Statistics table for extended figure panels 2.2, with the average response computed on a fixed [0-1] sec window instead of the optimal window (as in Figure 2 D, E, G, K, L, N).

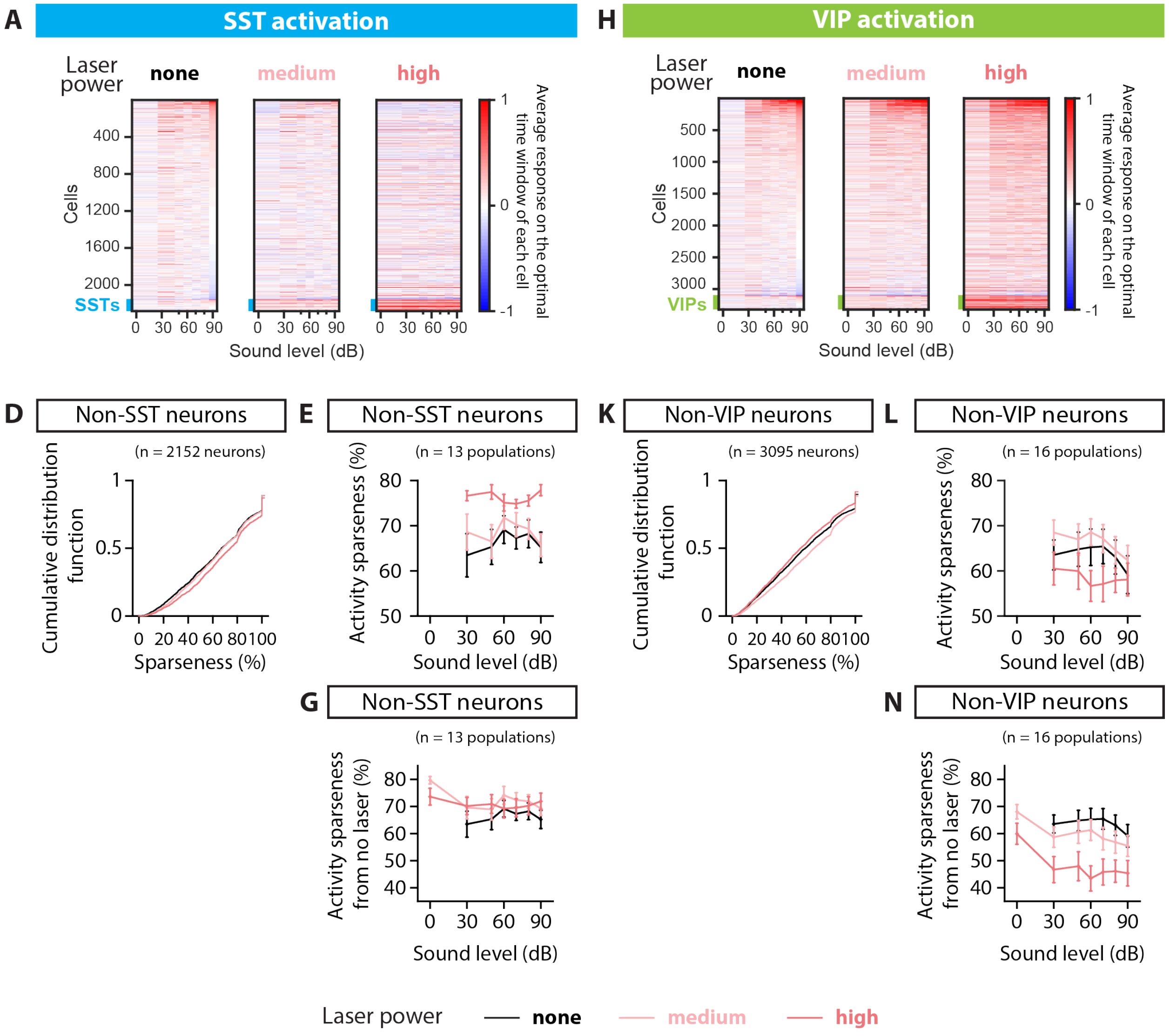


Extended Figure 2.2 : Panels A, D, E, G, H, K, L, N of Figure 2 computed on the fixed sec window as described in the Methods instead of the optimal window.

| **EXTENDED FIGURE 2.2** | | | | | | |
|---|---|---|---|---|---|---|
| Sparseness with SST activation | Ext. Fig 2.1 D | None: 2033<br>Med: 2029<br>High: 1994 | GLME | $t_{laser}$=5.21<br>DF = 5702 | *****$p_{laser}$=2.0e-7** | $h_{laser}^2$=4.5e-3 |
| Activity sparseness with SST activation | Ext. Fig 2.1 E | 13 populations | GLME | $t_{laser}$=3.20<br>$t_{sound}$=0.87<br>$t_{laser:sound}$=-0.79 | ****$p_{laser}$=1.6e-3**<br>$p_{sound}$=0.38<br>$p_{laser:sound}$=0.43 | $h_{laser}^2$=0.22<br>$h_{sound}^2$=1.3e-2<br>$h_{laser:sound}^2$=2.5e-2 |

| | | | | | | |
|---|---|---|---|---|---|---|
| | | | | DF = 230 | | |
| Activity sparseness from 0dB and no laser power with SST activation | Ext. Fig 2.1 G | 13 populations | GLME | $t_{laser}$=1.39<br>$t_{sound}$=0.84<br>$t_{laser:sound}$=-0.48<br><br>DF = 230 | $p_{laser}$=0.17<br>$p_{sound}$=0.40<br>$p_{laser:sound}$=0.63 | $h_{laser}^2$=3.9e-2<br>$h_{sound}^2$=9.0e-3<br>$h_{laser:sound}^2$=7.3e-3 |
| Sparseness with VIP activation | Ext. Fig 2.1 K | None: 2979<br>Med: 2921<br>High: 3020 | GLME | $t_{laser}$=-2.78<br><br>DF = 8350 | ****$p_{laser}$=5.5e-3** | $h_{laser}^2$=7.0e-4 |
| Activity sparseness with VIP activation | Ext. Fig 2.1 L | 16 populations | GLME | $t_{laser}$=-1.46<br>$t_{sound}$=-1.18<br>$t_{laser:sound}$=0.17<br><br>DF = 284 | $p_{laser}$=0.15<br>$p_{sound}$=0.24<br>$p_{laser:sound}$=0.86 | $h_{laser}^2$=2.8e-2<br>$h_{sound}^2$=1.2e-2<br>$h_{laser:sound}^2$=6.4e-4 |
| Activity sparseness from 0dB and no laser power with VIP activation | Ext. Fig 2.1 N | 16 populations | GLME | $t_{laser}$=-5.37<br>$t_{sound}$=-1.38<br>$t_{laser:sound}$=0.60<br><br>DF = 284 | *****$p_{laser}$=1.7e-7**<br>$p_{sound}$=0.17<br>$p_{laser:sound}$=0.55 | $h_{laser}^2$=0.19<br>$h_{sound}^2$=9.7e-3<br>$h_{laser:sound}^2$=4.6e-3 |

Extended Table 2.2: Statistics table for extended figure panels 2.1, with the average response computed on the fixed window as described in the Methods instead of the optimal window (as in Figure 2 D, E, G, K, L, N).
